# A sulfur-cycling co-culture alters iron sulfide diagenesis and pyritization

**DOI:** 10.64898/2026.10.20.753417

**Authors:** Kinga Sántha, Arnaud Duverger, Johanna Marin-Carbonne, Jasmine S. Berg

## Abstract

Iron sulfide minerals play a key role in modulating the oxygenation of Earth’s atmosphere and represent an archive for reconstructing past environmental conditions. In anoxic sedimentary environments, microbially produced sulfide reacts with iron, thereby forming metastable iron sulfides (mackinawite, greigite) which are expected to transform into pyrite, the most thermodynamically stable phase. However, our understanding of this process mostly stems from pure cultures grown on sulfate, which fails to capture the complexity of natural microbial communities and subsequent sedimentary changes. To this end, we set up pure and co-culture incubations of sulfate- and sulfur-reducing bacteria with selected ferric minerals (goethite, ferric phosphate and ferrihydrite) or ferrous iron and later subjected the samples to experimental diagenesis at controlled temperature (80 °C). We hypothesized that the co-culture would enhance pyrite formation by maintaining an internal cryptic S cycle that replenishes polysulfides and ferrous iron via the sulfidation of ferric compounds. Our findings showed that iron sulfide morphology was strongly dependent on the reactivity of ferric compounds, rather than the presence of bacteria. Consistent with this, pyrite formed through diagenesis in both pure cultures and abiotic incubations amended with sodium sulfide, indicating that pyritization is a primarily chemically controlled process. Contrary to our expectations, the presence of the co-culture led to longer diagenetic preservation of metastable minerals like greigite and marcasite, ultimately hindering pyritization. Surprisingly, we could also detect spherulitic pyrite after experimental diagenesis in sterile samples without exogenous sulfide addition. This was explained by the abiotic disproportionation of elemental S which took place in temperature regimes usually associated with microbial sulfide production. Overall, these results highlight the influence of mixed microbial communities, but also overlooked abiotic reactions, on iron sulfide (trans)formations in sedimentary environments, with implications for paleo(magnetic) reconstructions and the usability of these minerals as biosignatures.

## 1 Introduction

Iron sulfides are ubiquitous minerals derived from microbially produced sulfide in Earth’s surface environments (Picard et al., 2016). Pyrite, as the most stable and abundant phase, modulates the oxygenation of Earth’s atmosphere by its long-term burial (Berner, 1983; Halevy et al., 2012). Moreover, the isotopic composition of pyrite and trace elements co-precipitated with it are often used to reconstruct past environmental conditions (Mansor et al., 2025). Pyrite was also proposed to be a promising biosignature candidate, i.e. “phenomenon that provides evidence for the existence of life” (Catling et al., 2018; Runge et al., 2023), because its size, morphology, sulfur (S) and iron (Fe) isotopic composition and association with organic matter may bear distinctive characteristics when pyrite forms in the presence of microbes or microbially derived sulfide (Böttcher et al., 1998; Duverger et al., 2021; Marin-Carbonne et al., 2023; Nabeh et al., 2022; Truong et al., 2024). Pyrite precipitation occurs via three currently recognized reaction pathways: the hydrogen sulfide (H_2_S) pathway (Rickard, 1997; Rickard & Luther, 1997), polysulfide pathway (Luther, 1991; Rickard, 2012) and ferric-hydroxide-surface pathway (Peiffer et al., 2015; Wan et al., 2017). These reactions usually progress through a series of metastable iron sulfide intermediates such as mackinawite (FeS) and/or greigite (Fe_3_S_4_) (Hunger & Benning, 2007; Rickard & Luther, 2007). While these pathways and intermediates are independently well constrained in laboratory experiments focusing on abiotic reactions, they might be operating simultaneously in natural habitats. However, their prevalence and thus contribution to pyrite formation are likely dependent on environmental conditions (Rickard, 2012; Wan et al., 2017), many of which are affected by microbial activity.

Microbes have in fact been shown to further influence the formation of iron sulfides beyond the production of sulfide (Schoonen, 2004). For example, it was suggested that microbes can locally increase the concentration of sulfide and/or polysulfides at the surface of ferric minerals (Berg et al., 2020) or elemental S_(0)_ (Sekerci et al., 2025), and thus accelerate reaction rates (Picard et al., 2018). Furthermore, negatively charged cellular surfaces were proposed to act as mineral nucleation sites (Konhauser, 1998). Iron sulfides precipitated in the presence of microbes were often found to exhibit crystallographic differences from their abiotic counterparts such as larger size and tendency to aggregate (Picard et al., 2018); and formation of distinct associations with organic matter (Picard et al., 2021; Truong et al., 2023). Nevertheless, many previous experiments utilizing sulfate-reducing bacteria and ferrous iron managed to precipitate only mackinawite and greigite (Benning et al., 1999; Fortin et al., 1994; Gramp et al., 2010; Herbert et al., 1998; Picard et al., 2018; Williams et al., 2005; Zhou et al., 2014), even though pyrite could have formed during the reaction of FeS and sulfide (H_2_S-pathway) (Picard et al., 2018). However, bacterial biomineralization experiments supplied with ferric compounds (goethite, ferric phosphate) or different S sources (elemental S_(0)_ instead of sulfate) precipitated pyrite (Canfield et al., 1998; Duverger et al., 2020; Finster et al., 1998; Rickard, 1969; Sekerci et al., 2025). In these cases, pyrite formation is thought to be mediated by polysulfides, which form in the reaction of sulfide with elemental S_(0)_ following the reductive dissolution of ferric compounds (Duverger et al., 2020; Rickard, 1969; Sekerci et al., 2025). Polysulfides may also be produced in the oxidation of sulfide by O_2_ or manganese oxides, highlighting their prevalence in natural habitats (Avetisyan et al., 2021; Chen & Gupta, 1973).

In sum, existing laboratory studies focusing on Fe-S biomineralization have mostly grown pure cultures in the presence of one type of S compound. By contrast, iron sulfides in natural environments form in a complex interplay between microbially and abiotically mediated reactions which produce a series of S compounds with oxidation states between sulfate and sulfide. S cycle intermediates such as elemental S_(0)_, thiosulfate and tetrathionate often do not accumulate to high concentrations in natural habitats (Zopfi et al., 2004), and have therefore often been neglected in laboratory incubations. However, recent studies point toward their rapid microbial recycling in cryptic cycles (Findlay & Kamyshny, 2017; Vigneron et al., 2021), thus potentially contributing to a supply of sulfide for mineral precipitation. Lastly, once formed, iron sulfides continue to experience further physico-chemical changes as the sediment layers are buried deeper during diagenesis (Aller, 2014; Berner, 1983). These transformations are caused by an increase in temperature and pressure, the effects of which are often overlooked in laboratory incubations where iron sulfides are precipitated at atmospheric pressure and temperature optimal for bacterial growth (Duverger et al., 2020; Picard et al., 2018; Sekerci et al., 2025).

To address these gaps, we investigated iron sulfide formation and transformation in the presence of a co-culture containing the sulfate-reducing bacterium *Desulfovibrio desulfuricans* and elemental S_(0)_-reducing bacterium *Sulfurospirillum deleyianum*. These bacteria were previously shown to co-occur and lead to the rapid precipitation of pyrite in an enrichment culture prepared from the water of ferruginous lake Pavin (Berg et al., 2020; Hansel et al., 2015). We hypothesized that sulfide production by these bacteria in the presence of different ferric iron minerals would initiate an internal cryptic S cycle (Fig. 1). The continuous reductive dissolution of Fe(III) would constantly replenish ferrous iron and S_(0)_, thus promoting pyritization (Mansor et al., 2025). After a period of intense bacterial sulfide production, we also subjected the (bio)minerals to experimental diagenesis to investigate the effects of temperature increase during sediment burial. The potential role of co-cultures and/or experimental diagenesis in biomineral formation was assessed by comparing iron sulfide morphology and mineralogy to those formed in pure cultures or abiotic incubations externally amended with sodium sulfide. The morphological and mineralogical results presented here expand our knowledge of iron sulfide mineral (trans)formations and preservation in systems more closely resembling the complexity of natural environments, with implications for their usability as paleoindicators or biosignatures on modern or ancient Earth alike.

**Figure 1.**
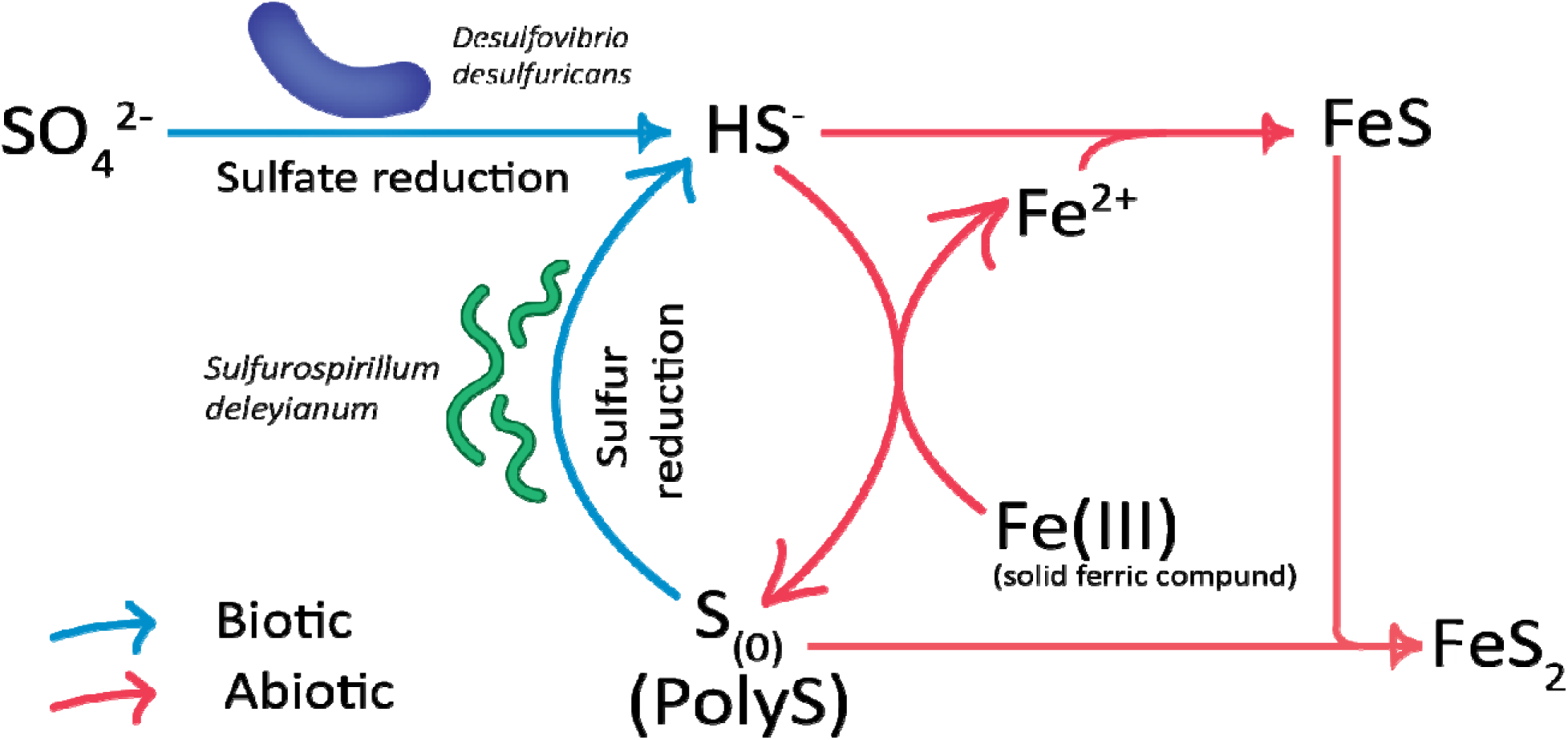
Schematic of the (hypothetical) internal cryptic S cycle mediated by *D. desulfuricans* and *S. deleyianum*. Abbreviations: PolyS – polysulfides.

## 2 Material and methods

### 2.1 Cultivation and routine maintenance of bacteria

Viable cultures of the model bacteria *Desulfovibrio desulfuricans* (DSM 642, subsequently *D. desulfovibrio*) and *Sulfurospirillum deleyianum* (DSM 6946, subsequently *S. deleyianum*) are regularly maintained at “name of institution”. The growth media are composed of the following solutions: salt solution for *D. desulfuricans* (Duverger et al., 2020), salt solution for *S. deleyianum* (Straub & Schink, 2004), 1 mL L^-1^ trace element solution S4178 from MediaDive (Koblitz et al., 2023), 1 mL L^-1^ selenite/tungstate solution (Tschech & Pfennig, 1984), 10 mL L^-1^ 1X Wolin’s vitamin solution S5980 from MediaDive (Koblitz et al., 2023), 3.00 g L^-1^ MOPS buffer, with electron donors and acceptors added separately. In the case of *D. desulfuricans*, 10 mM sodium sulfate and 20 mM sodium DL-lactate are used as electron acceptor and donor, respectively. *S. deleyianum* is fed 25 mM sodium fumarate, 10 mM sodium formate, and 5 mM sodium acetate as an additional carbon source to fumarate, plus 0.5 mM L-cysteine hydrochloride monohydrate as a S source. After mixing the salts, trace element, selenite/tungstate and electron donor/acceptor solutions, the pH is adjusted to 7-7.2 with 1 M NaOH and the medium is dispensed into 100 mL serum vials. The serum vials are hermetically sealed with butyl rubber septa and aluminium caps, heated at 55 °C for several minutes, degassed with 100 % N_2_ gas and autoclaved (20 min at 121 °C and 2 atm). The sterile and anoxic vitamin solution is added prior to inoculating 4 % v/v bacterial suspension into a total of 50 mL growth medium. The bacteria are incubated at 30 °C in the dark without shaking for several days. Later, *D. desulfuricans* is transferred to room temperature (19-20 °C) to slow down the poisoning caused by hydrogen sulfide. Bacteria are transferred into fresh media (4% v/v inoculum) every 4-6 weeks.

### 2.2 Preparation of the biomineralization media

Four iron compounds were chosen for the setup of the biomineralization experiments based on their abundance in nature and tendency to form pyrite in previous experiments (Duverger et al., 2020; Picard et al., 2016; Rickard, 1969; Sun et al., 2018): ferrous iron (as FeCl_2_), ferrihydrite (Fe_10_O_14_(OH)_2_.nH_2_O), ferric phosphate (FePO_4_) and goethite (α-FeO(OH)). Ferrous iron was prepared as a 1 M solution of FeCl_2_ * 4 H_2_O (referred to as ferrous iron or Fe^2+^ in later text) and sterile filtered in a Jacomex glovebox (N_2_ atmosphere, O_2_ < 5 ppm).

Goethite powder was commercially purchased (Sigma-Aldrich, 71063), while nanoparticulate ferrihydrite and ferric phosphate were synthesized in-house as outlined in Cornell & Schwertmann (2003) and Mirvaux et al. (2016), respectively. The powders were stored in the dark.

Each of the four iron minerals were tested under the following conditions: three biotic incubations (a pure culture of *D. desulfuricans*, a pure culture of *S. deleyianum* and a co-culture of the two bacteria), one sterile abiotic incubation and one sterile control. In the abiotic incubations, a sterile solution of sodium sulfide was manually injected to simulate bacterial sulfide production. No sodium sulfide was added to the controls to check for potential abiotic transformation of the iron compounds. Biotic incubations were performed in triplicate, whereas abiotic incubations and controls were not replicated due to time and space constraints. An overview of the experiments is provided in Table 1. The color-coding of Table 1 corresponds to the exact medium composition outlined in Table 2.

**Table 1.**
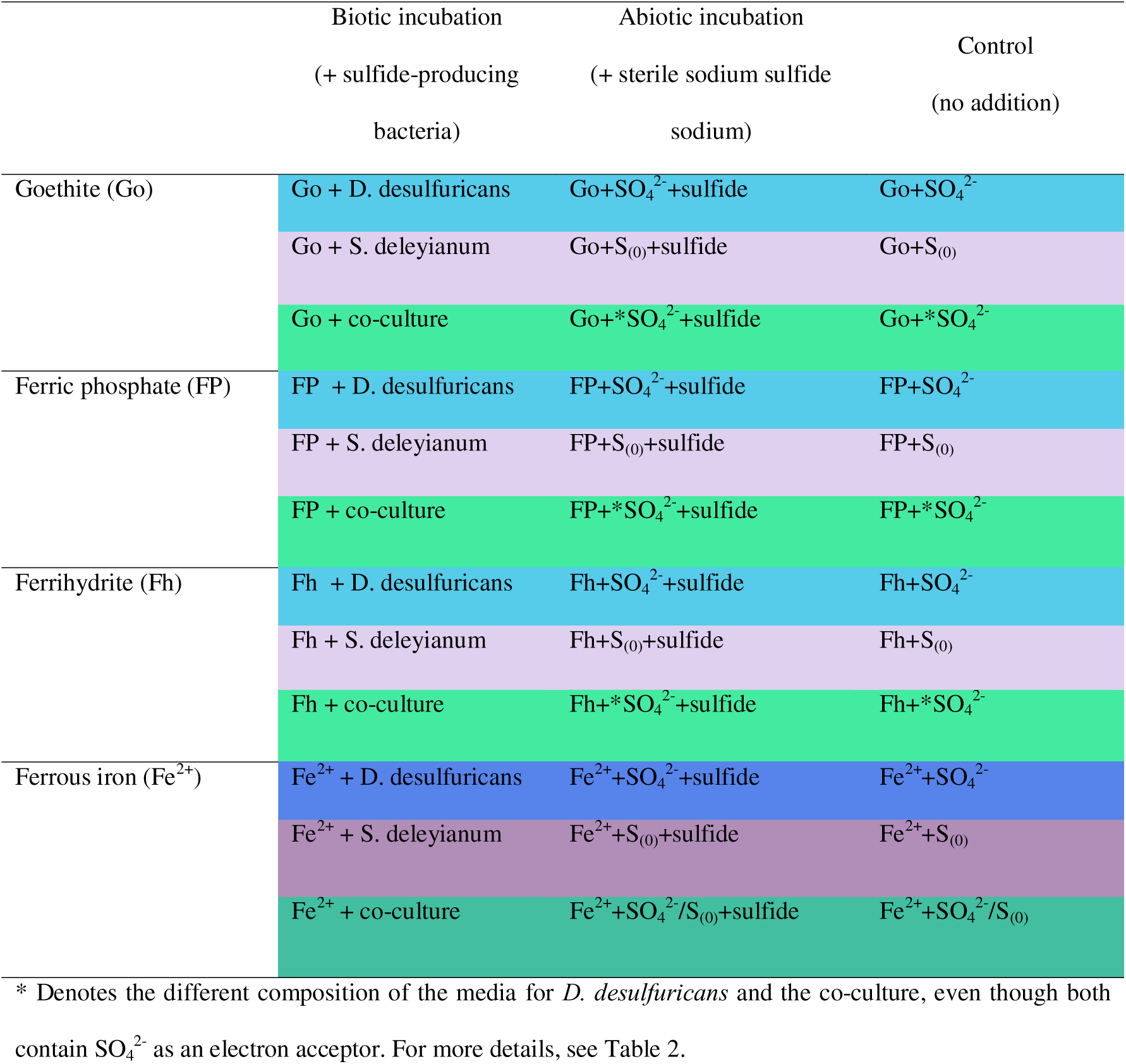
Summary of the biomineralization experiments with four different types of added iron. For each biotic incubation, an abiotic incubation with added sodium sulfide and a control incubation with no additions was prepared using the same medium composition, denoted by incubation names. Biotic incubations were performed in triplicate. The color-coding corresponds to the concentration and combination of electron donors/acceptors used in the respective experiments, outlined in more detail in Table 2.

**Table 2.**
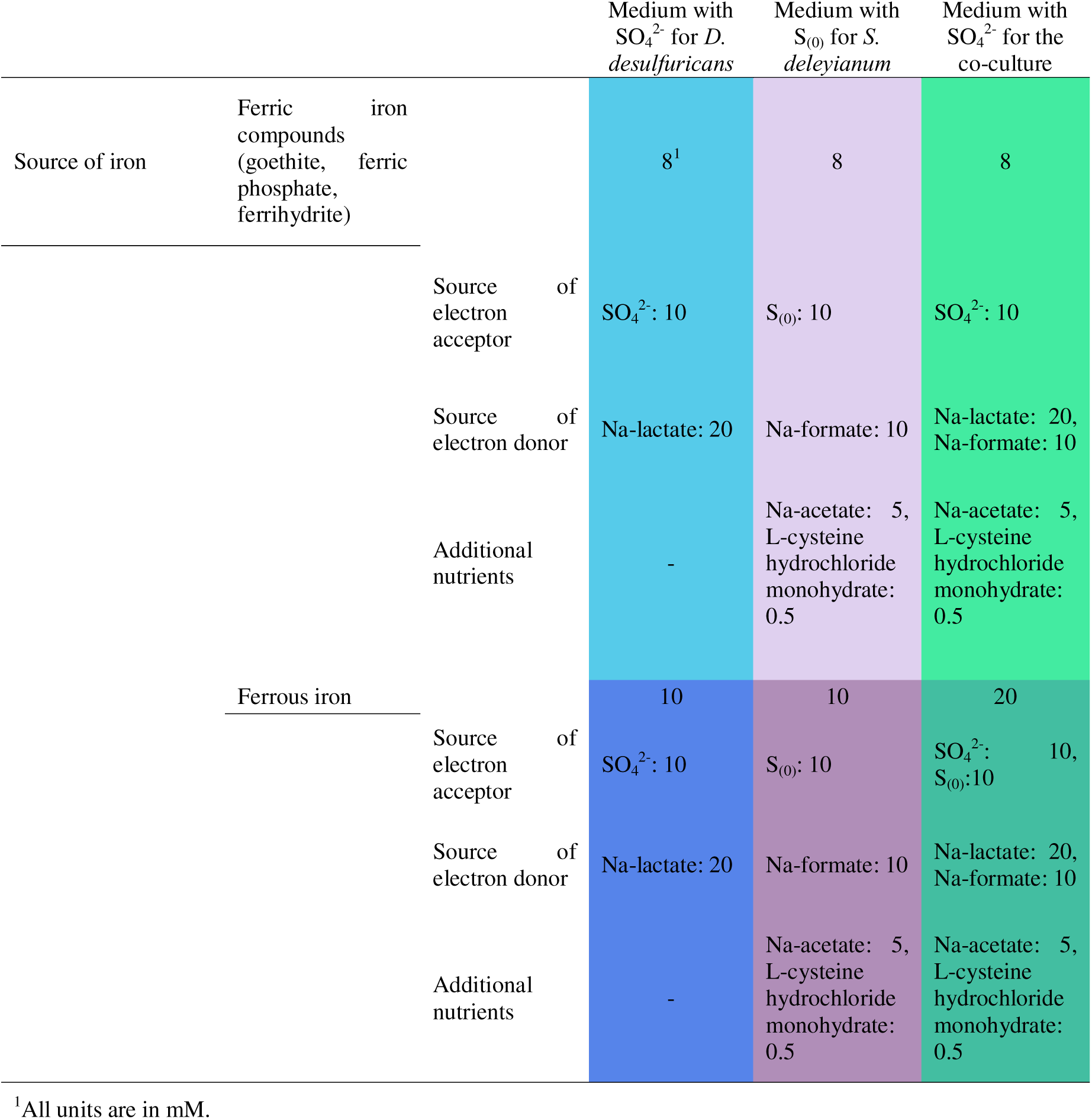
Composition of the media used in the four biomineralization experiments (goethite, ferric phosphate,

|  |  |  | Medium with<br>SO<sub>4</sub><sup>2-</sup> for *D.*<br>*desulfuricans* | Medium with<br>S<sub>(0)</sub> for *S.*<br>*deleyianum* | Medium with<br>SO<sub>4</sub><sup>2-</sup> for the<br>co-culture |
| --- | --- | --- | --- | --- | --- |
| Source of iron | Ferric iron<br>compounds<br>(goethite, ferric<br>phosphate,<br>ferrihydrite) |  | 8<sup>1</sup> | 8 | 8 |
|  |  | Source of<br>electron<br>acceptor | SO<sub>4</sub><sup>2-</sup>: 10 | S<sub>(0)</sub>: 10 | SO<sub>4</sub><sup>2-</sup>: 10 |
|  |  | Source of<br>electron donor | Na-lactate: 20 | Na-formate: 10 | Na-lactate: 20,<br>Na-formate: 10 |
|  |  | Additional<br>nutrients | - | Na-acetate: 5,<br>L-cysteine<br>hydrochloride<br>monohydrate:<br>0.5 | Na-acetate: 5,<br>L-cysteine<br>hydrochloride<br>monohydrate:<br>0.5 |
|  | Ferrous iron |  | 10 | 10 | 20 |
|  |  | Source of<br>electron<br>acceptor | SO<sub>4</sub><sup>2-</sup>: 10 | S<sub>(0)</sub>: 10 | SO<sub>4</sub><sup>2-</sup>: 10,<br>S<sub>(0)</sub>: 10 |
|  |  | Source of<br>electron donor | Na-lactate: 20 | Na-formate: 10 | Na-lactate: 20,<br>Na-formate: 10 |
|  |  | Additional<br>nutrients | - | Na-acetate: 5,<br>L-cysteine<br>hydrochloride<br>monohydrate:<br>0.5 | Na-acetate: 5,<br>L-cysteine<br>hydrochloride<br>monohydrate:<br>0.5 |

<sup>1</sup>All units are in mM.

The biomineralization media were prepared fresh, i.e. maximum one week before the start of the experiments. The liquid components of the medium (salt solution for *D. desulfuricans*, trace element, selenite/tungstate solutions and the electron donors/acceptors) were mixed and sterilized as described above. The ferric iron minerals were sterilized by tyndallization (heated at 90 °C three times for 1 h on three consecutive days) in 100-mL serum vials. Their crystal structures were not significantly affected by heating as verified by XRD (Fig S1). Similarly, the S_(0)_ powder was tyndallized in microcentrifuge tubes (heated at 90 °C three times for 3 h on three consecutive days). The S_(0)_ powder, the liquid medium and the vitamin solution were added to the serum vials containing the solid iron minerals in a laminar hood, followed by aseptic degassing with N_2_. For the ferrous iron biomineralization experiment, the FeCl_2_ * 4 H_2_O solution was added in the glovebox after combining the rest of the medium. The final volume of medium in each replicate was 50 mL with a 50 mL N_2_ headspace. The final pH of the buffered medium before bacterial inoculation was 7-7.2 and it was not further monitored during the incubation period. All serum vials were protected from light during storage.

### 2.3 Setup of the biomineralization experiments

The setup of the biomineralization experiments, subsequent subsampling and sample preparation were carried out under an inert (N_2_) atmosphere in a Jacomex^®^ glovebox. *S. deleyianum* and *D. desulfuricans* were cultured in growth media (outlined in Section 2.1) for 1 and 9 days, respectively, until mid- to late logarithmic phase. Growth was monitored by measuring protein concentrations following the method of Bradford as outlined in Ceriotti et al. (2025). As growth curves made by measuring protein concentrations possessed the characteristic logarithmic and stationary phases, protein concentration was found to be a reliable proxy for tracking growth changes over time (Ceriotti et al., 2025). The bacterial cultures were centrifuged at 7000 g for 10 min and washed three times with 15-25 mL of sterile and anoxic salt solution for *D. desulfuricans*. After the last washing, the bacterial pellets were resuspended in 5 mL salt solution and added to the biomineralization media using sterile syringes and needles in equal mass amounts, as calculated by the protein concentrations. The final concentration of bacterial suspensions in 50 ml medium varied from 6.5 to 10 ug/mL protein between the different biomineralization experiments. Regarding the available electron acceptors, the medium where *D. desulfuricans* was inoculated contained only sodium sulfate, while the medium for *S. deleyinaum* contained only S(0). The medium for the co-cultures contained sodium sulfate, while S_(0)_ was expected to form internally by the sulfidation of the ferric compounds (see Tables 1 and 2). In the case of the abiotic incubations and controls, 5 mL sterile and anoxic salt solution (used for the preparation of the medium) was added instead of bacteria. Furthermore, abiotic incubations were supplemented with a sterile solution of 1 M Na_2_S * 9 H_2_O in a final concentration matching that of electron acceptors (sodium sulfate and/or S_(0)_).

Immediately after the setup (T_0_), subsamples of 5 mL were aseptically taken for solid mineral analysis (except in the ferrous iron experiment where there was no solid precipitate in the bacterial cultures or control). Serum vials were placed in an incubator at 30 °C in the dark without shaking for four weeks, which allowed for intense bacterial sulfide production. After this incubation period, 20 mL suspension was subsampled from each vial. The remaining solution in the serum vials was transferred to an 80 °C oven to simulate experimental diagenesis (Mansor et al., 2025; Višić et al., 2025). This temperature was much higher than the optimal growth temperature of the bacteria and placed the diagenetic conditions at ∼ 2.5-3 km depth given a geothermal gradient of 25-30 °C km^-1^. Diagenesis was simulated for 15-16 weeks, which allowed for the prolonged exposure of minerals to heat. After the diagenetic incubation, a second subsampling followed. The subsamples were pelleted by centrifugation at 7000 g for 15 min and washed 3x by resuspension in anoxic MQ and centrifugation for 5 min. The samples were left to dry in microcentrifuge tubes and stored anoxically in the glovebox. Before downstream analysis, triplicate samples from the biotic incubations were pooled to form a composite sample.

### 2.4 Morphological characterization and identification of the iron sulfide precipitates

#### 2.4.1 Scanning electron microscopy (SEM)

A part of the dried powder samples was resuspended in 8-10 mL anoxic MQ inside the glovebox and filtered onto 13 mm polycarbonate GTTP (pore size 0.2 um) filters using Swinnex filter holder units. The polycarbonate filters were deposited onto adhesive carbon tape on aluminium stubs and sputter-coated with carbon. The samples were imaged using a Zeiss GeminiSEM 460 (for the T_0_ samples) and Zeiss GeminiSEM 500 scanning electron microscopes, both equipped with a field emission gun (FEG) and an Oxford AZtec Microanalysis System (Oxford Instruments, High Wycombe, UK).

The surface characteristics of the minerals were observed with the help of secondary electrons generated by a high-performance annular detector at low voltage conditions (5 kV) and at 3-5 mm working distance. Backscattered electron images (generated by an angle selective backscattered detector), as well as elemental characterization by energy dispersive X-ray spectrometry (EDS), were acquired at 8-10 mm working distances and 20 kV voltage. The images taken were further cropped in Adobe Illustrator 27.9.6.

#### 2.4.2. Transmission electron microscopy (TEM)

Only bacterial incubations from the goethite experiment prior to diagenesis were analyzed to assess the presence of minerals that could have remained undetected by XRD or Raman spectroscopy (see below). Part of the dried powder samples were resuspended in anoxic ultrapure water until turbidity was no longer visible to the naked eye (approximately 1-2 mL). Three microliters of this suspension were deposited onto the 200-mesh copper TEM grids coated with a carbon support film, then left to dry in the glovebox overnight. The TEM grids were observed at the IMPMC TEM plateform under a JEOL 2100F transmission electron microscope equipped with a field emission gun (FEG) operating at 200 kV. Scanning transmission electron microscopy (STEM) was performed in the high-angle annular dark field (HAADF) mode and coupled to energy dispersive X-ray spectrometry (EDS) mapping. Energy-dispersive X-ray spectra were processed from EDS datacube by Hyperspy Python library (Peña et al., 2020) available for download at https://github.com/hyperspy/hyperspy. Additionally, selected-area electron diffraction (SAED) patterns were acquired on areas of interest.

#### 2.4.3 X-ray diffraction (XRD)

The dried powder samples were deposited onto an air sensitive sample holder (Bruker) and anoxically sealed inside the glovebox. Diffraction patterns were acquired at 40 kV and 40 mA with Cu-Kα radiation (λ=0.154 nm) on a ThermoFisher ARL Equinox 1000 diffractometer. The measurements lasted for 1-1.5 hours. The acquired diffractograms were further analyzed in the Match! software (Crystal Impact, version 4.1). The baseline was automatically detected and subtracted by Match! and all diffractograms were moved 0.2-0.6 ° to account for the variable sample position during measurements. The peaks were annotated using the Crystallography Open Database (COD).

#### 2.4.4 Confocal Raman spectroscopy

Powder samples were resuspended in anoxic MQ and inserted into rectangular capillary tubes (5 cm * 2 mm, CM Scientific) inside the glovebox. The ends of capillary tubes were glued onto glass slides (5 cm * 7.5 cm) with wax to minimize O_2_ diffusion and prevent drying. Alternatively, 20-40 uL of resuspended sample was deposited onto the glass slide and covered with a microscope cover slip which was sealed with electric tape to prevent the diffusion of O_2_ into the sample. All samples were analyzed shortly after being prepared.

Raman spectra were recorded using a Renishaw inVia Qontor spectrometer coupled to a confocal Leica microscope, with a 532 nm laser source using a 1800 l mm^-1^ grating and 50-100X magnification. Approximately 3-10 recordings were made in each sample with 1-5 mW laser power and 1-10 accumulations. The baseline was automatically detected and subtracted by the WiRE™ software (5.6). The recorded spectra were identified using the in-built spectrum search option of WiRE, the RRUFF database (https://rruff.info/), Bourdoiseau et al. (2008) for (oxidized and aged) mackinawite, Chi Fru et al. (2018) for pyrite and marcasite, and Eder et al. (2014) for greigite and magnetite, Mirabello et al. (2021) for ferric phosphate, Dzieniszewska et al. (2020) for ferrihydrite, and Liu et al. (2017) for (ferrous) phosphate minerals.

### 2.5 (Follow-up) experiments for abiotic reactions of Fe and S at elevated temperatures (80 °C)

In the absence of sulfide-producing bacteria or externally added sulfide, black precipitates appeared in the abiotic controls with elemental S_(0)_ and Fe^2+^+SO_4_^2-^after experimental diagenesis. To verify our results, follow-up experiments were conducted. The media were prepared as outlined in Section 2.2 under strictly sterile and anoxic conditions. To study the effect of heating on elemental S_(0)_, two sets of triplicate vials were incubated at 30 and 80 °C for up to 8 months. Liquid phase subsampling took place at the start of the experiment (T_0_), after 1 and 16 weeks. The subsamples (1.5-2 mL) were taken with N_2_-flushed sterile syringes inside the laminar flow hood, filtered on 0.22 µm PTFE filters and stored frozen until further analysis. Samples for mineralogy were analyzed after 8 months as outlined above.

To investigate the effect of heating sulfate in the presence of ferrous chloride, two rounds of experiments were run in triplicates. In the first round, one set of triplicate vials were supplemented with both 10 mM sodium sulfate and 10 mM ferrous chloride, while the other included only 10 mM sodium sulfate. The second round was set up in a similar way except the addition of sodium sulfate, i.e. it contained a set of triplicate vials with only 10 mM ferrous chloride and a set with only sterile medium without ferrous chloride or sodium sulfate. Subsamples were taken in the glovebox at the start of the experiment (T_0_), four weeks at 30 °C and 16 weeks of heating at 80 °C. Liquid phase subsamples were collected as mentioned above. Subsamples for dissolved iron - Fe_D_ (1 mL; assumed to be composed mostly of ferrous iron) were centrifuged and the supernatant was preserved in 1 mL anoxic 1 M HCl. At the end of the heating period, the rest of the solid phase samples were preserved for mineralogical analyses as described in previous sections. The summary of the follow-up experiments can be seen in Suppl. Table 1.

Liquid phase subsamples were acidified with a drop of 1 M HCl, degassed for 3 min with N_2_ gas to evacuate any sulfide present and diluted 10 times in MQ water. The concentrations of sulfate, thiosulfate and organic acids (lactate, formate) were measured with ion chromatography (Metrohm, 930 Compact IC Flex). Dissolved iron was quantified using ICP-OES (Agilent 5900) after a tenfold dilution in the sample matrix (0.5 M HCl).

## 3 Results

A detailed mineralogical and morphological summary of all biomineralization experiments is available in the Supplementary Material (Suppl. Tables 2-5).

### 3.1 Morphology and identity of iron sulfide minerals formed in goethite biomineralization experiments

Goethite was detected as short acicular crystals (Fig. S2) with well-defined peaks in the X-ray diffractograms (Fig. S3) at the start of the incubation (T_0_). After four weeks of incubation at 30 °C, a stark contrast was observed between the biotic and other incubations. While goethite remained unchanged in the abiotic incubations and controls (Fig. 2D-I), bacterial sulfide production led to the formation of aggregates of nanoparticulate iron sulfides (Fig. 2A-C; EDS-data not shown) coupled to the diminishment of goethite’s characteristic XRD peaks (Fig. S3). Oxidized (Fe^3+^-containing) mackinawite could be detected by Raman spectroscopy in all biotic incubations (Fig. 3A) (Bourdoiseau et al., 2008; Runge et al., 2024). Greigite was clearly identified in *S. deleyianum* cultures, whereas the occasional presence of a Raman band at 351 cm^-1^ (Eder et al., 2014) might also be attributed to its presence in *D. desulfuricans* and co-culture incubations. While pyrite could not be detected with XRD or Raman spectroscopy, TEM analyses (Fig. S4) revealed that it was present as a minor phase in co-cultures.

**Figure 2.**
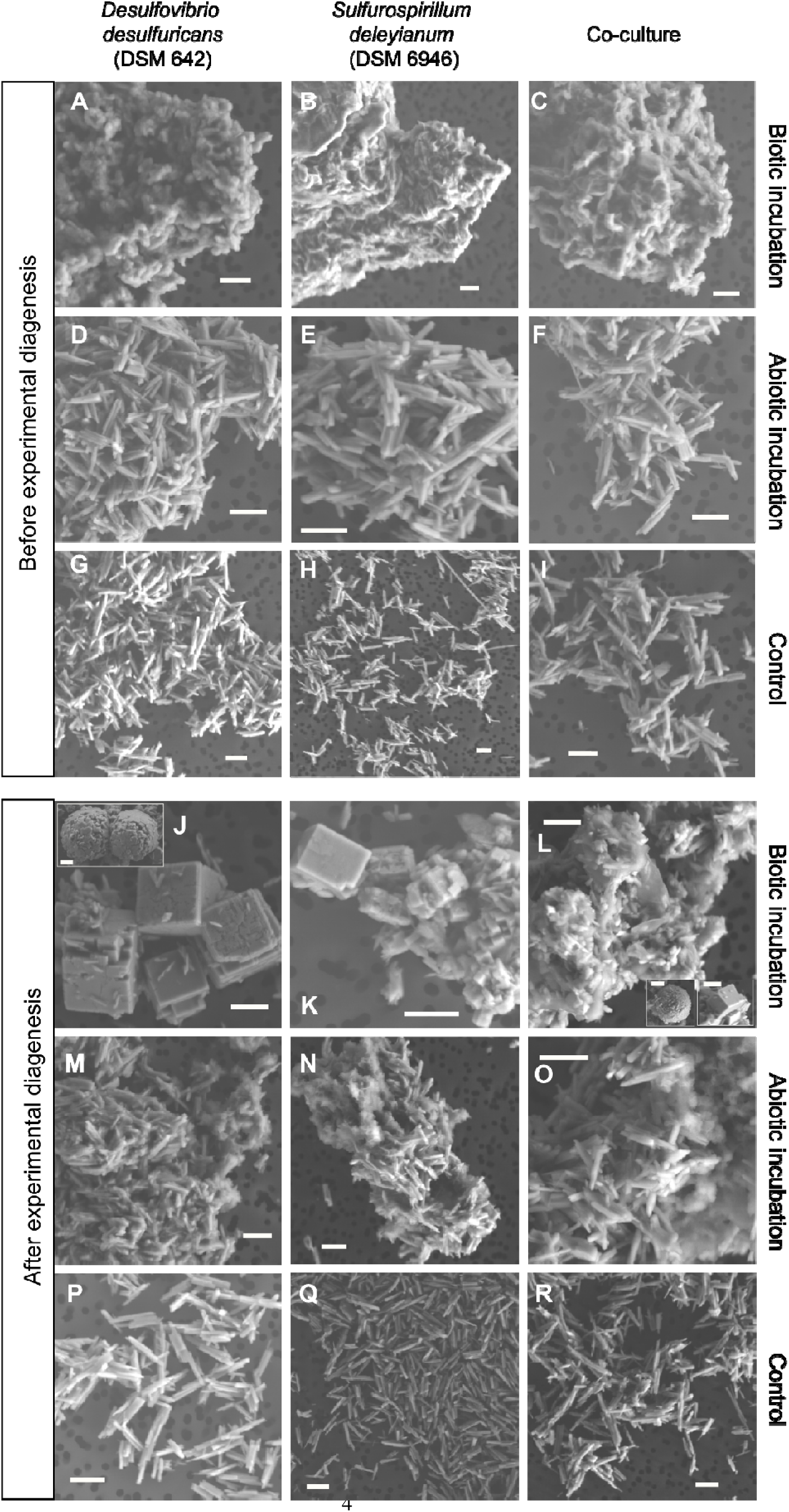
SEM images of the goethite biomineralization experiment. Scalebar is 1 um. All images were taken in secondary electron (SE) mode.

**Figure 3.**
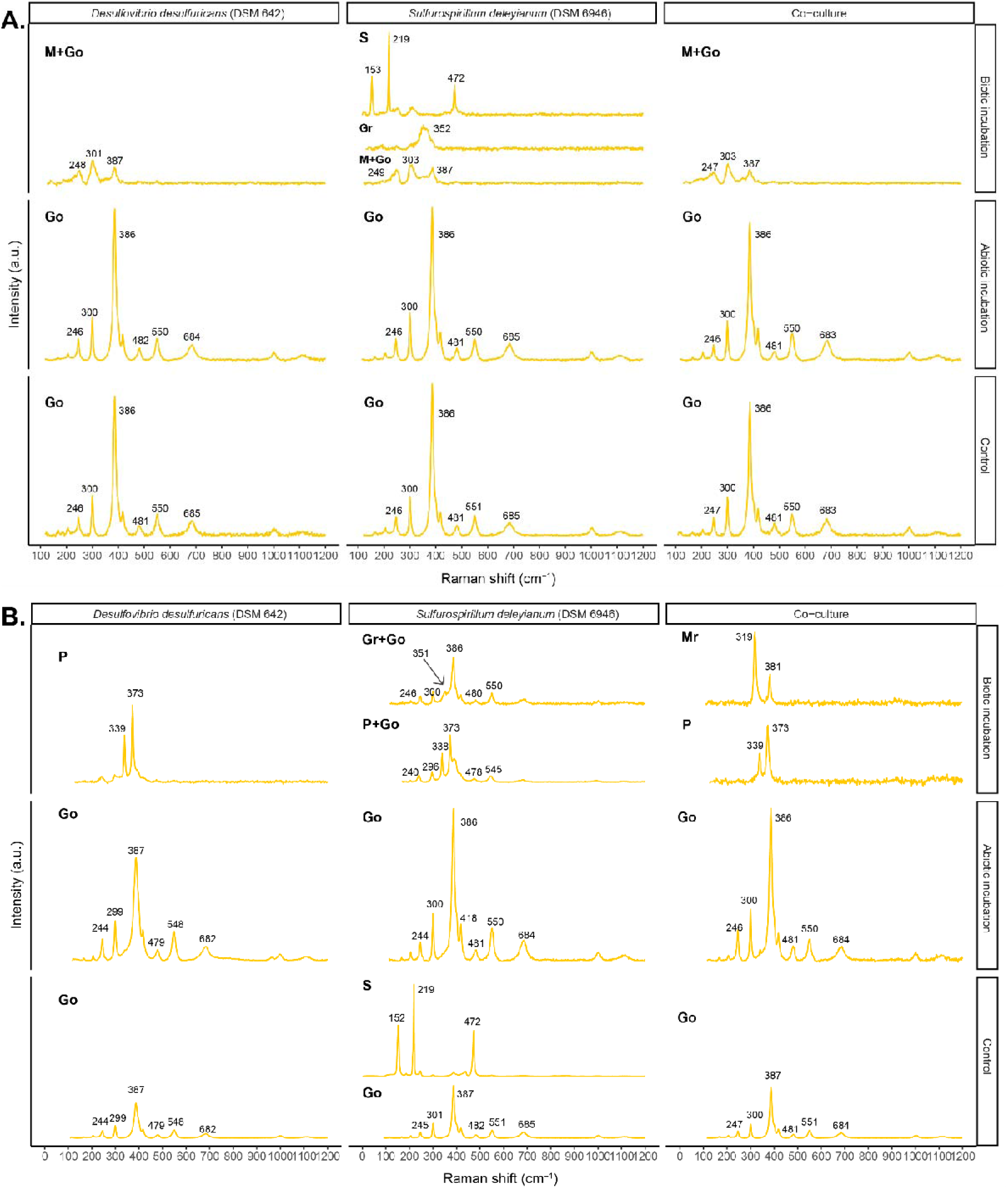
Raman spectroscopy results of the goethite biomineralization experiment. A. Incubations after 1 month at 30 °C. B - Incubations after 15-16 weeks of experimental diagenesis (80 °C). Abbreviations: P – pyrite, Mr – marcasite, M – mackinawite, Gr – greigite, S – elemental S_(0)_, Go – goethite.

After 16 weeks of experimental diagenesis at 80 °C, goethite in the abiotic incubations and controls retained its initial morphology (Fig. 2M-R). In contrast, the morphology of iron sulfides in the bacterial cultures was greatly affected (Fig. 2J-L). Namely, a gradual change in shape and decrease in individual grain size were observed which were most pronounced in the co-cultures. Micrometric euhedral (cubic) iron sulfides formed in the presence of *D. desulfuricans* cultures (Fig 2J). These minerals could be found individually or in aggregates. In some instances, submicrometric euhedra were closely packed to each other and assumed an overall spherical shape, resembling framboid-like structures. Besides the euhedral habit, micrometric spherules could also be detected. These minerals were identified as pyrite by XRD (Fig. S3) and Raman spectroscopy analyses (Fig. 3B). The remains of goethite (interspersed between the pyrite grains as short needles) were identified as well.

Iron sulfides from *S. deleyianum* cultures after diagenesis consisted of cube-like or irregularly shaped (sub)micrometric crystals that formed aggregates (Fig. 2K). In co-culture samples, iron sulfides appeared to be even smaller than with *S. deleyianum* and resembled irregularly shaped crystals that formed aggregates with goethite needles. In both these incubations, individual micrometric crystals shaped as cubic euhedra or spherules were sometimes still visible. The change in crystal morphology was concurrent with the appearance of multiple iron sulfides. In addition to pyrite, we could identify greigite and marcasite in the *S. deleyianum* and co-culture samples, respectively (Fig. 3B).

### 3.2 Morphology and identity of iron sulfide minerals formed in ferric phosphate biomineralization experiments

Ferric phosphate resembled colloform aggregates of fine spherical particles at the start of the experiment (T_0_, Fig S5). After four weeks at 30 °C, it remained untransformed in the controls (Fig. 4G-I). In contrast, the sulfidation of ferric phosphate occurred in both biotic and abiotic incubations, as indicated by the appearance of iron sulfide aggregates composed of irregularly shaped particles (Fig. 4A-F, EDS-data not shown). In *S. deleyianum* cultures, two types of morphologies were observed: (1) particles with well-defined edges containing P, O and Fe (EDS, not shown), or (2) aggregates of (nano)particles rich in Fe and S (Fig. 4B). Oxidized (Fe^3+^-containing) mackinawite and pyrite were detected by Raman spectroscopy in both pure cultures, as well as all abiotic incubations (Fig. 5A). However, in co-culture samples, we could identify only oxidized and aged mackinawite after four weeks (Fig. 5A). It cannot be excluded that pyrite was also present here, but its amount and/or degree of crystallinity might have been insufficient for detection by our methods.

**Figure 4.**
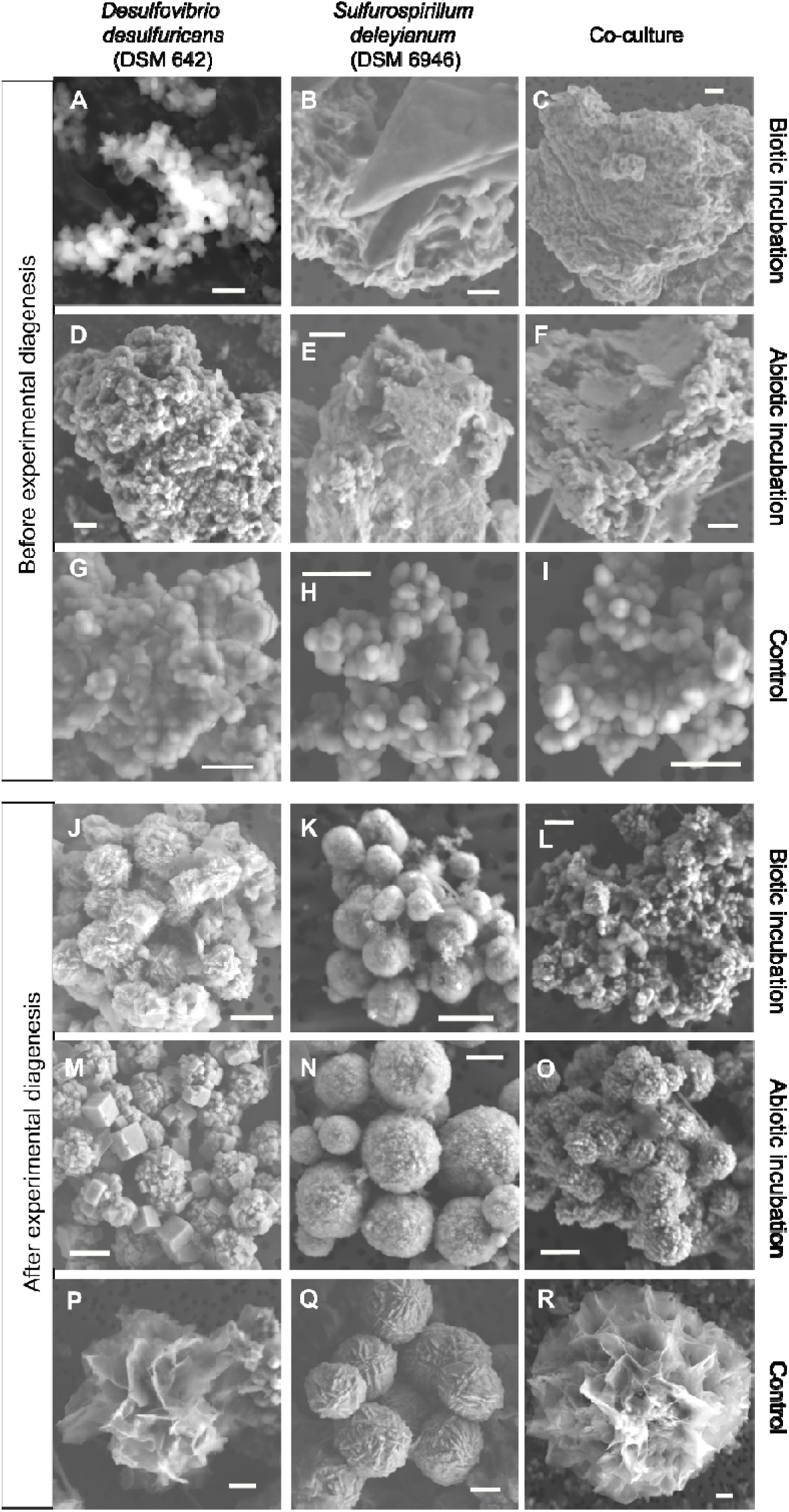
SEM images of the ferric phosphate biomineralization experiment. Scalebar is 1 um. All images, except A, were taken in secondary electron (SE) mode. Image A was made using backscattered electrons (BSD).

**Figure 5.**
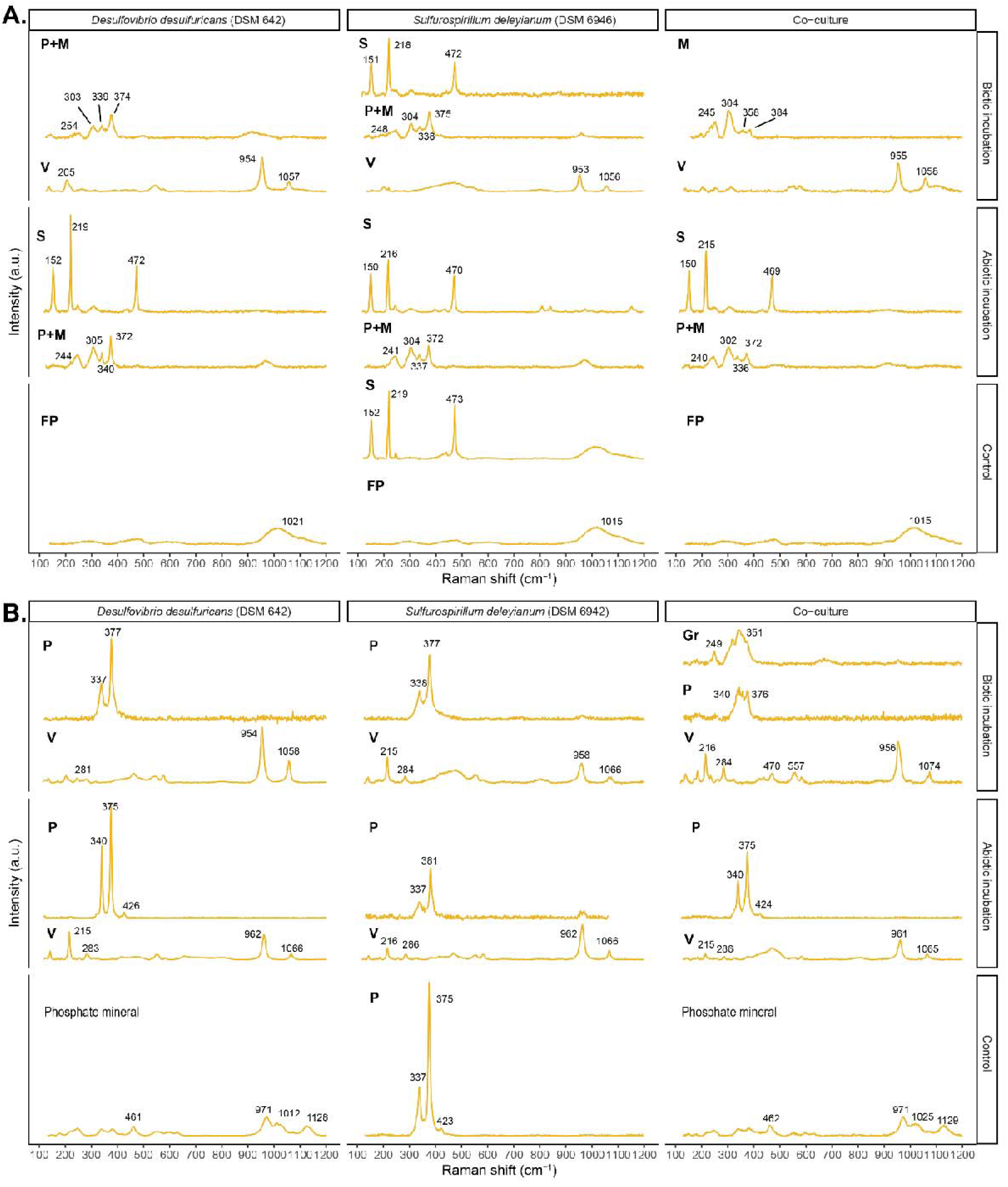
Raman spectroscopy results of the ferric phosphate biomineralization experiment. A. Incubations after 1 month at 30 °C. B - Incubations after 15-16 weeks of experimental diagenesis (80 °C). Abbreviations and explanations: P – pyrite, M – mackinawite, Gr – greigite, S – elemental S_(0)_, V – vivianite, FP – ferric phosphate, Phosphate mineral – non-vivianite-like phosphate mineral.

Simulated diagenesis led to profound changes in all samples. Structures of flower-like morphology appeared in all controls (Fig. 4P-R). These structures contained P, Fe and O (EDS-data not shown); identified as non-vivianite phosphate minerals (Fig. 5B). Surprisingly, the control with FP+S_(0)_ turned faintly black compared to the other controls. The visual change in color was later explained by the presence of (sub)micrometric spherulitic clusters rich in Fe and S (Fig. 4Q). With the aid of both XRD and Raman spectroscopy, the identity of these spherules was confirmed to be pyrite (Fig. 5B, S6).

Micrometer-sized spherules containing Fe and S were found in *D. desulfuricans* (Fig. 4J) and *S. deleyianum* cultures (Fig. 4K); as well as in the abiotic incubations containing FP+S_(0)_+sulfide and FP+*SO_4_^2-^+sulfide (Fig. 4N-O) after diagenesis. These spherules commonly formed aggregates. Micrometric euhedral (cubic) iron sulfides appeared in the FP+SO_4_^2-^+sulfide incubation (Fig. 4M). Aggregates of submicrometric cubic-shaped crystals were also visible, resembling framboid-like structures. The identity of these minerals was confirmed to be pyrite (Fig. 5B, S6). In co-culture samples, iron sulfides decreased in size as compared to pure cultures, similar to co-cultures incubated in the presence of goethite. In more detail, submicrometric irregularly shaped or cube-like minerals formed aggregates (Fig. 4L), composed mostly of greigite and, tentatively, pyrite (Fig. 5B, S6).

### 3.3 Morphology and identity of iron sulfide minerals formed in ferrihydrite biomineralization experiments

After four weeks at 30 °C, ferrihydrite retained its poorly defined morphology (Fig. S7) in the controls (Fig. 6G-I). In contrast, aggregates of (nano)particulate minerals appeared in biotic and abiotic incubations (Figs. 6B-F). These structures contained O, Fe and S in variable amounts (EDS point spectra, not shown), which could be due to the incomplete sulfidation of ferrihydrite. In co-culture samples, two shapes could be distinguished: (1) aggregates composed of submicrometric spherules and (2) sheets (Fig. 6C) like those commonly attributed to mackinawite (Csákberényi-Malasics et al., 2012; Picard et al., 2018; Runge et al., 2024). The presence of oxidized (Fe^3+^-containing) mackinawite was further corroborated in all biotic and abiotic incubations with the help of Raman spectroscopy (Fig. 7A). Unfortunately, attempts at cultivating *D. desulfuricans* in the presence of ferrihydrite were unsuccessful.

**Figure 6.**
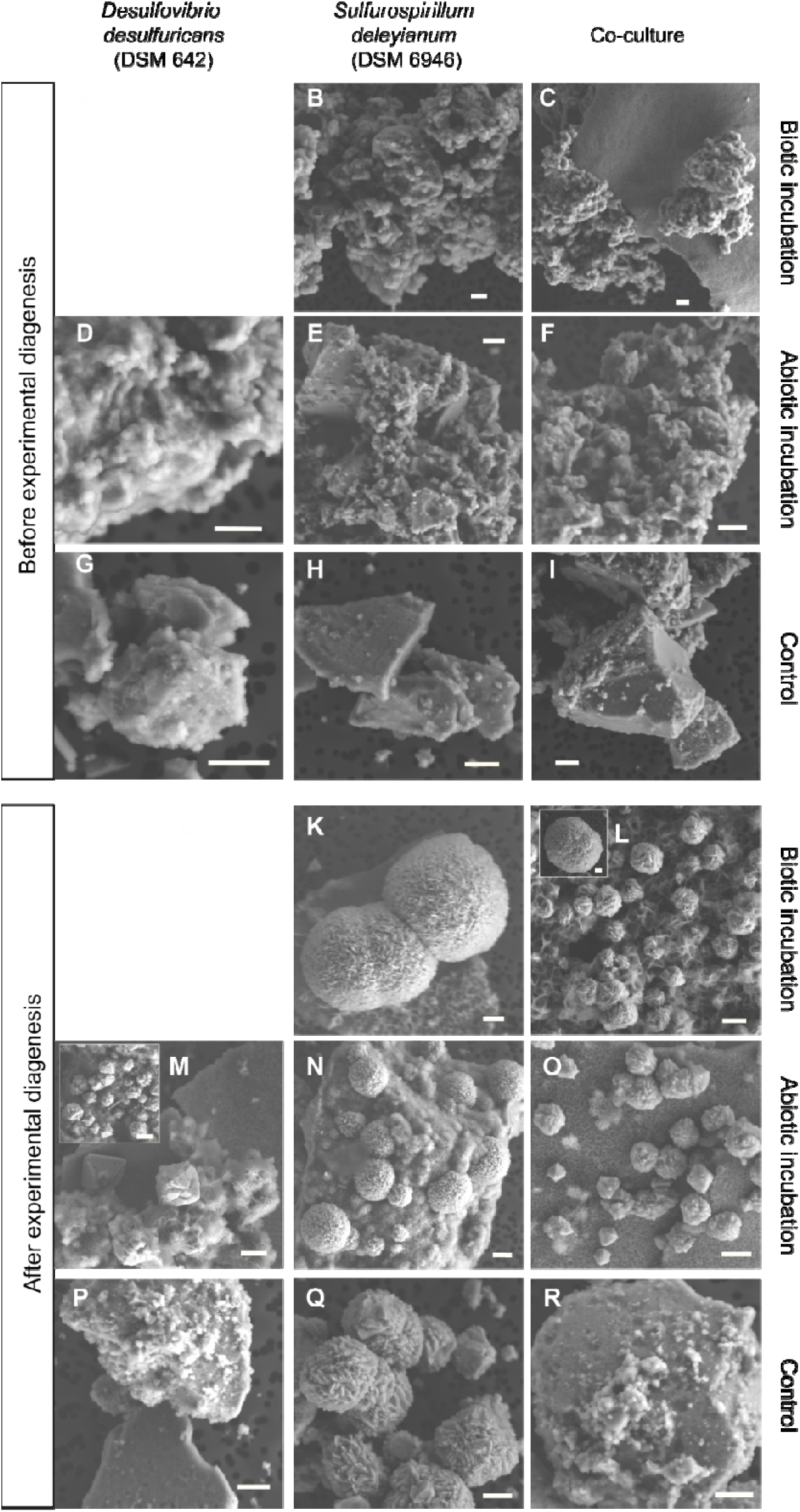
SEM images of the ferrihydrite biomineralization experiment. Scalebar is 1 um. All images were taken in secondary electron (SE) mode.

**Figure 7.**
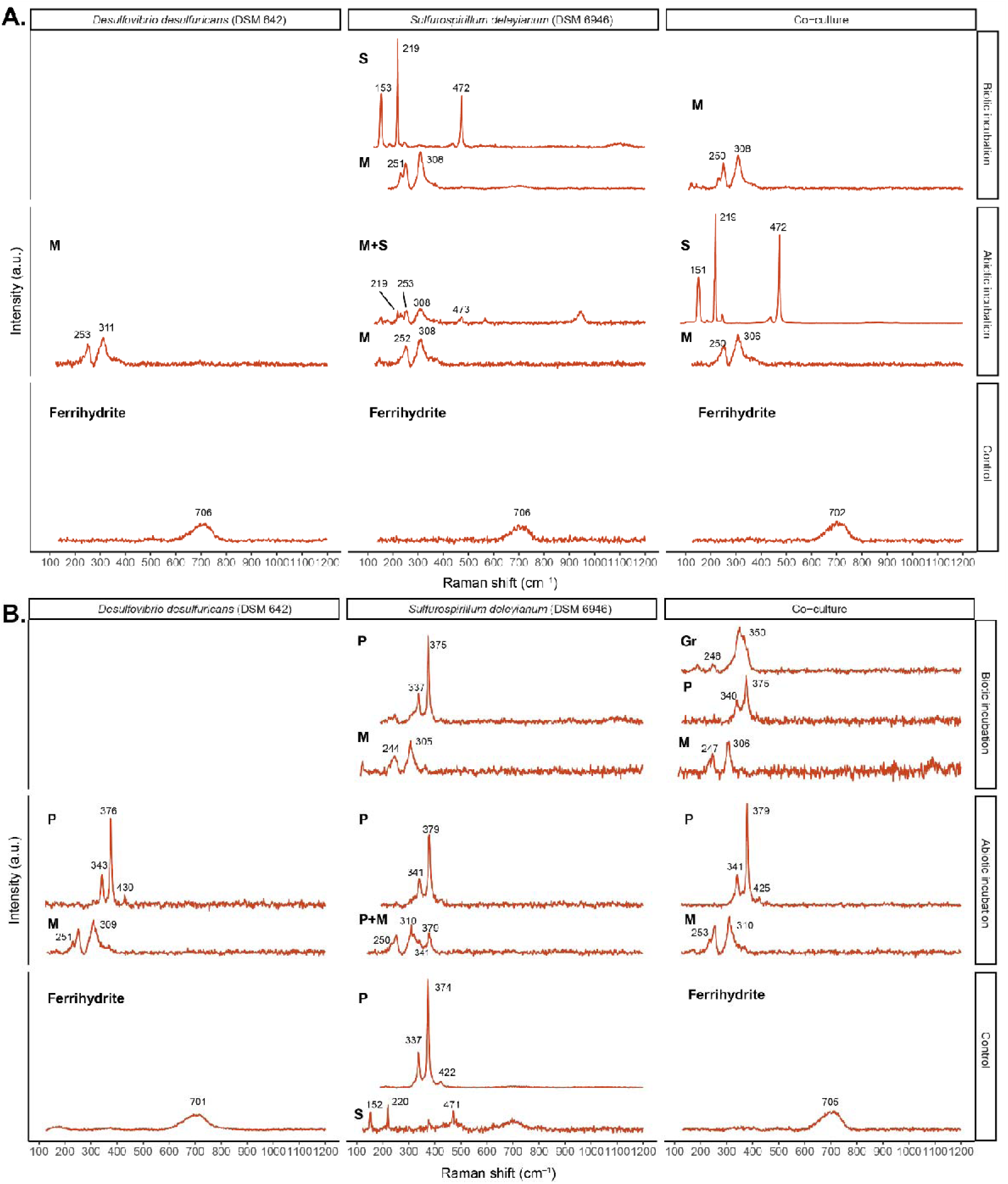
Raman spectroscopy results of the ferrihydrite biomineralization experiment. A. Incubations after 1 month at 30 °C. B - Incubations after 15-16 weeks of experimental diagenesis (80 °C). Abbreviations: P – pyrite, M – mackinawite, Gr – greigite, S – elemental S_(0)_.

Experimental diagenesis led to profound transformations in almost all samples containing ferrihydrite except for controls with Fh+SO_4_^2-^ and Fh+*SO_4_^2-^ (Fig. 6P, R). The control with Fh+S_(0)_ turned gray and contained spherulitic clusters of Fe- and S-rich minerals (Fig. 6Q), similar to incubations with ferric phosphate. These minerals were confirmed to be pyrite (Fig. 7B, S8).

Sheet-like and micrometric spherulitic iron sulfides appeared in all biotic and abiotic incubations (Fig. 6K-O) after diagenesis. Moreover, the abiotic incubations with Fh+SO_4_^2-^+sulfide and Fh+*SO_4_^2-^+sulfide contained micrometric euhedral (octahedral) crystals as well (Fig. 6M, O). The identity of these minerals was confirmed to be pyrite and oxidized (Fe^3+^-containing) mackinawite (Fig. 7B, S8). Given that pyrite belongs to the isometric crystal system, the euhedra and spherules can be attributed to this mineral, whereas mackinawite likely appeared as sheets. Besides pyrite and mackinawite, greigite was also identified in the presence of co-cultures (Fig. 7B, S8), like in co-cultures with ferric phosphate. However, in contrast to incubations with goethite and ferric phosphate, no concurrent change in mineral shape or decrease in size was observed after experimental diagenesis.

### 3.4 Morphology and identity of iron sulfide minerals formed in ferrous iron biomineralization experiments

After four weeks at 30 °C, iron sulfide aggregates composed of (nano)particles were visible in all incubations with added ferrous iron (Fig. 8A-F). These minerals were mostly identified as mackinawite (Fig. 9A, S9). In more detail, mackinawite was detected in its reduced form in *D. desulfuricans* cultures and Fe^2+^+SO_4_^2-^+sulfide incubation (Fig. 9A); while the oxidized form was present in other samples. Greigite could also be identified in *S. deleyianum* cultures by the presence of a single broad Raman band at ∼ 350 cm^-1^ (Fig. 9A). The presence of this peak, although diminished, might indicate greigite in other samples as well: *D. desulfuricans* culture, co-culture, and abiotic incubations with Fe^2+^+S_(0)_+sulfide and Fe^2+^+*SO_4_^2-^+sulfide. Alternatively, the co-occurrence of smaller peaks at ∼350-360 and 385-386 cm^-1^ with the characteristic mackinawite peak at ∼300-305 cm^-1^ might suggest the presence of aged mackinawite, similar to the co-cultures in the presence of ferric phosphate.

**Figure 8.**
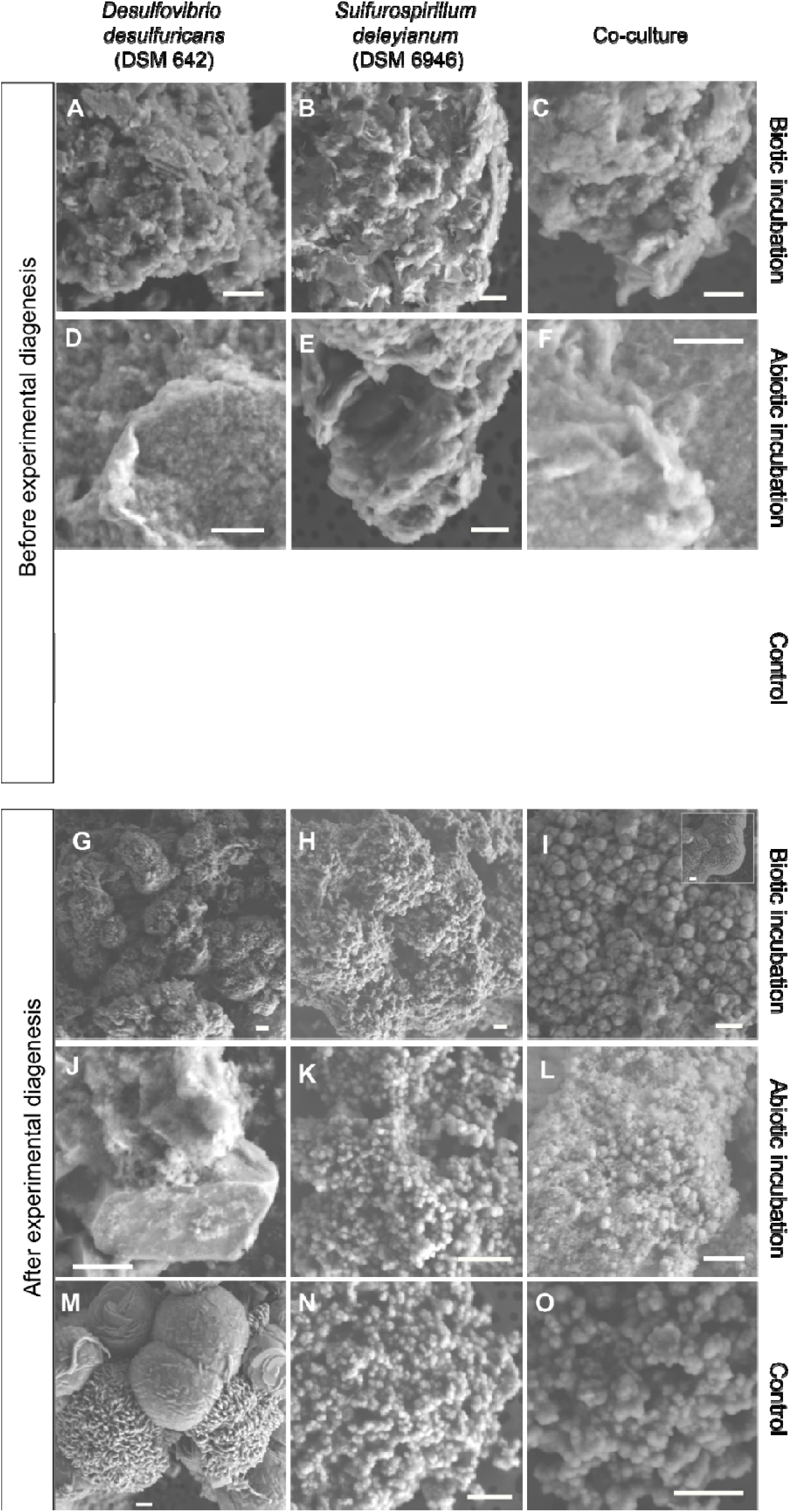
SEM images of the ferrous chloride biomineralization experiment. Scalebar is 1 um. All images were taken in secondary electron (SE) mode.

**Figure 9.**
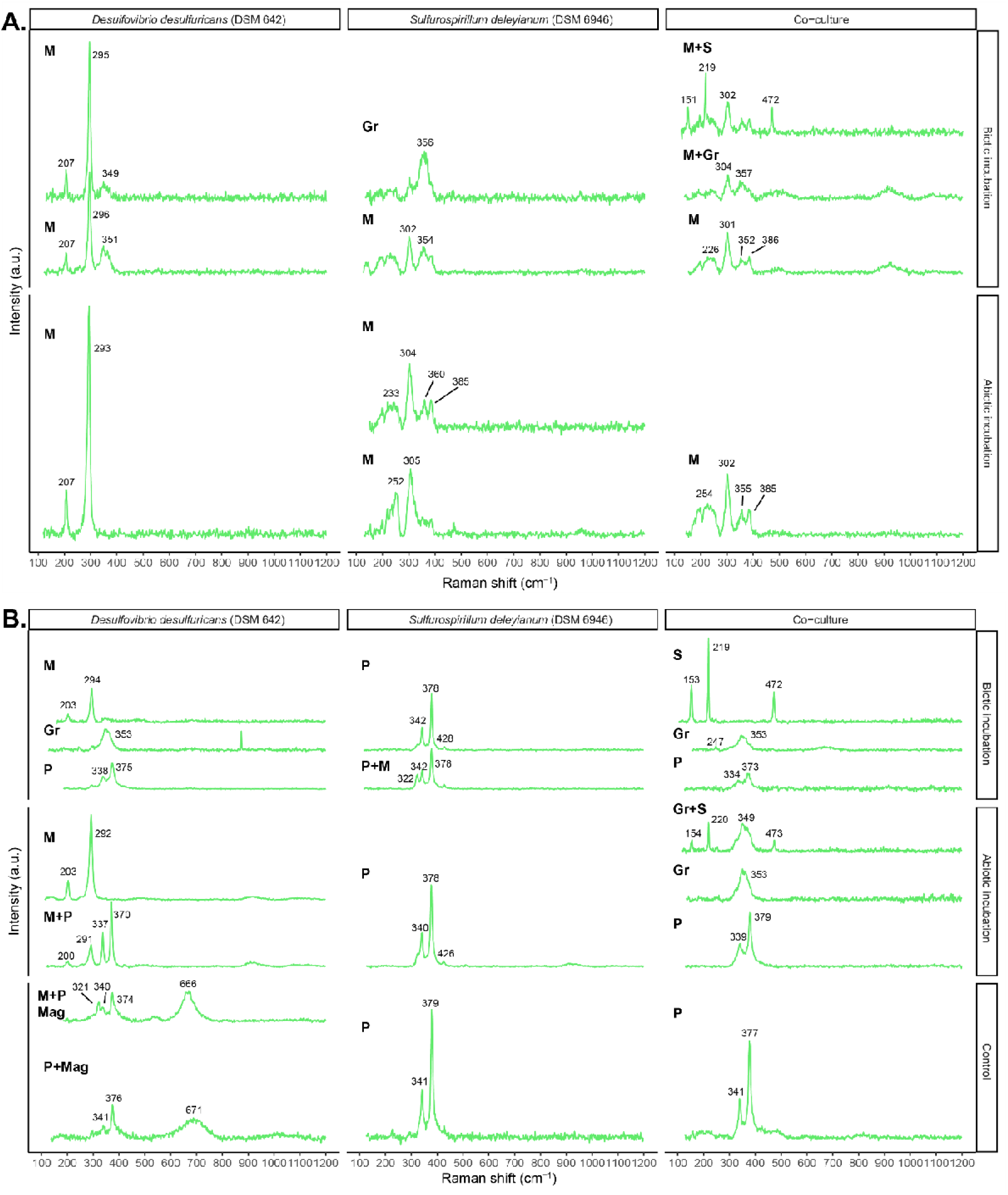
Raman spectroscopy results of the ferrous iron biomineralization experiment. A. Incubations after 1 month at 30 °C. B -Incubations after 15-16 weeks of experimental diagenesis (80 °C). Abbreviations: P – pyrite, M – mackinawite, Gr – greigite, S – elemental S_(0)_, Mag – magnetite.

Following 16 weeks of experimental diagenesis, the controls turned black in all setups. The control with Fe^2+^+SO_4_^2-^contained spherules (Fig. 8M) and (sub)micrometric octahedral crystals. These structures were rich in Fe and S (EDS, not shown) and confirmed to be pyrite (Fig. 9B, S9). In addition to characteristic Raman peaks of pyrite, a peak at ∼ 320 cm^-1^ could be attributed to oxidized (Fe^3+^-containing) mackinawite (Bourdoiseau et al., 2011; Runge et al., 2024) and several others to magnetite. All controls containing Fe^2+^+S_(0)_ harbored aggregates of submicrometric spherules (Fig. 8N-O) of pyrite (Fig. 9B, S9).

The heated abiotic incubations with Fe^2+^+SO_4_^2-^+sulfide contained iron sulfides of irregular shape with the occasional appearance of euhedral crystals identified as reduced mackinawite and pyrite, respectively (Fig. 8J, 9B, S9). In the abiotic incubation with Fe^2+^+S_(0)_+sulfide and Fe^2+^+SO_4_^2-^/S_(0)_+sulfide, submicrometric spherules composed of pyrite and a combination pyrite and greigite, respectively, were identified (Fig. 8K-L, 9B, S9). Despite the presence of S_(0)_ in both these incubations, the transformation of greigite to pyrite was not complete in all cases. Since greigite is a known precursor in pyrite formation (Hunger & Benning, 2007), we hypothesize that the physical proximity of these two phases in sample Fe^2+^+SO_4_^2-^/S_(0)_+sulfide (Fig. 8L) points toward an example of slow pyritization, as was observed previously (Kubeneck et al., 2025).

The experimental diagenesis of pure cultures produced iron sulfide (nano)particles aggregated into larger spherules (Fig. 8G-H). These were composed of a combination of mackinawite, greigite and pyrite in *D. desulfuricans* cultures (Fig. 9B, S9). Cultures of *S. deleyianum* contained pyrite and, occasionally, oxidized mackinawite (peak at ∼ 322 cm^-1^). The co-cultures contained (1) submicrometric spherules that formed aggregates and (2) larger spherules (Fig. 8I). Greigite seemed to dominate these samples based on the obtained well-defined XRD peaks (Fig. S9), similar to co-cultures with other iron sources after simulated diagenesis. Although not visible in bulk XRD analysis probably due to abundance levels below the detection limit, pyrite could be detected in Raman point spectra (Fig. 9B).

### 3.5 Abiotic reactions of Fe and S at elevated temperatures (80 °C)

#### 3.5.1 Transformation of elemental S_(0)_ under experimental diagenetic conditions

All controls supplemented with elemental S_(0)_ powder (FP+S_(0)_, Fh+S_(0)_, Fe^2+^+S_(0)_ and Fe^2+^+SO_4_^2-^/S_(0)_) turned dark gray after experimental diagenesis which was later explained by the presence of spherulitic pyrite. To verify whether elemental S_(0)_ could be the source of sulfide at this temperature, we set up additional sterile experiments containing elemental S_(0)_ and ferrihydrite. At the start of the incubation period, all vials were the dark red color of ferrihydrite when shaken (Fig. S10A). After only one week, samples incubated at 80 °C turned dark (gray to black), whereas those at 30 °C retained their initial color (Fig. S10B). No further visual changes could be observed until the end of the experiment (Fig. S10C). The change in coloration was concurrent with an increase in thiosulfate concentrations over time, reaching ∼ 1 mM at the end of the incubation period (Fig. S10H-I). On the other hand, no increase in sulfate concentrations was measurable. The concentration of formate remained stable over time. Mineralogical analyses revealed the presence of (sub)micrometric, S and Fe-rich spherules (Fig. S10D-G) which were shown to be pyrite (Fig. S10J-K).

#### 3.5.2 Abiotic S transformations in the presence of ferrous chloride under experimental diagenetic conditions

Due to the surprising formation of spherulitic pyrites in control experiments containing ferrous iron and sulfate after experimental diagenesis, follow-up experiments were performed. Sterile incubations were set up with different combinations of sulfate and ferrous chloride and subjected to 30 °C and 80 °C (see Suppl. Table 1: sulfate+ferrous iron, only sulfate, only ferrous iron, medium without sulfate or ferrous iron). After experimental diagenesis, bottles that did not contain ferrous iron remained clear, whereas ferrous iron-supplemented bottles turned dark irrespective of the presence of sulfate (Fig. S11B and S12C). The change in color was accompanied by a decrease in dissolved ferrous iron (from ∼9.2 to 5 mM), corroborating its partitioning into solid phase particles (Fig. S11I-J), while sulfate concentrations remained constant over time (∼ 10 mM). SEM analysis of the ferrous iron-containing samples revealed microstructures that contained Fe, S and O in variable combinations (EDS, not shown). Namely, sheets and (sub)micrometric spherules (Fig. S11C-E, S12D-E) were observed that mostly contained Fe and O. Moreover, spherules and plates rich in Fe and S were also visible (Fig. S11F-G, S12F-G). Mineralogical analyses confirmed the presence of pyrite and marcasite, but also magnetite and oxidized mackinawite (Fig. S11K-L, S12H-I), which contain ferric iron.

## 4 Discussion

### 4.1 The reactivity of ferric iron compounds influences the morphology of iron sulfide minerals

The extensive sulfidation of iron compounds in most experimental conditions led to the formation of morphologically similar aggregates prior to diagenesis and regardless of the presence of viable microbial populations (biotic and abiotic incubations before exp. diagenesis in Fig. 4, 6 and 8). This contrasts with previous work which found that biogenic minerals exhibited larger size, distinct morphologies and tendency to aggregate as opposed to abiogenic iron sulfides (Picard et al., 2018). In our study the relative size of mineral aggregates was directly evaluated by visually comparing SEM images, while Picard et al. (2018) calculated crystallite size based on the results of XRD or dynamic light scattering. This raises the possibility that subtle differences on the level of crystallite size might have remained unnoticed when visually observing larger mineral aggregates in our samples. In addition, our experiments were significantly shorter than previous work, so we cannot exclude the potential evolution of the observed minerals if left over longer periods of time (Duverger et al., 2020; Gramp et al., 2010; Li et al., 2006; Picard et al., 2018; Sekerci et al., 2025).

The only clear visual differences between biotic and abiotic incubations before diagenesis occurred in the goethite biomineralization experiment (Fig. 2A-F). As opposed to abiotic incubations where the initial goethite morphology remained unchanged in the presence of sterile sodium sulfide, microbial sulfidation produced nanoparticulate aggregates of iron sulfides. The sulfidation of goethite was shown to be accelerated by low(er) pH (Rickard, 1974). Actively metabolizing bacteria can create local low-pH microniches inside biofilms despite buffering of the bulk medium (Hidalgo et al., 2009; Hwang et al., 2016; Nguyen et al., 2022; Xiao et al., 2017). As our cultures were static during the entire incubation period, it is likely that bacterial aggregates locally acidified the pH on goethite surfaces despite the bulk medium being buffered at ∼ pH 7-7.2. Consequently, Fe^3+^ on the surface of goethite could have been more easily solubilized, resulting in its subsequent reaction with sulfide (Hoffmann et al., 2021; Rickard, 1974). Furthermore, any number of other factors related to the presence of bacteria, such as direct enzymatic reduction (Li et al., 2006) or secretion of exopolymeric substances (EPS), might have further facilitated the dissolution of goethite. Consistent with this interpretation, in a similar study design Li et al. (2006) reported that 7.5 % of the starting goethite reacted with sodium sulfide, compared to 60-75 % in the presence of sulfate-reducing bacteria. In sum, these findings clearly highlight the influence of microbial activity on the sulfidation of certain iron oxides with high crystallinity and low solubility (e.g. goethite).

Incubation at 80 °C is expected to either significantly decrease bacterial metabolic rates or lead to cellular death (Duverger et al., 2021). Thus, we can assume that any new mineral precipitation during experimental diagenesis is chemically controlled, with microbes only playing a passive role as nucleation sites or as organic matter sources. The very similar mineral habits in pure cultures and abiotic incubations after diagenesis therefore suggest that the presence of dead bacterial biomass continues to exert a minor influence on morphology. Instead, morphology seems to change from euhedral (cubic) in goethite incubations to spherulitic pyrite in samples with ferrihydrite and ferric phosphate, reflecting changes in ferric compound reactivity (Figs. 2, 4 and 6). Ferric compound reactivity is known to control sulfidation rates (Canfield, 1989; Raiswell, 1982), i.e. the rate at which sulfide dissolves ferric compounds into ferrous iron and intermediate S compounds. Thus, the sulfidation kinetics of ferric compounds likely controlled the rate at which reactants needed for pyrite formation were supplied to the system, thereby generating different supersaturation levels with respect to pyrite. Previous studies demonstrated that pyrite adopts varying habits depending on temperature and supersaturation (Murowchick & Barnes, 1987; Rickard, 2012; Wang & Morse, 1996). Goethite is thus expected to react slowly with sulfide, as observed here and in previous studies (Lentini et al., 2012; Li et al., 2006). The gradual reaction of sulfide with goethite before and during experimental diagenesis would generate a steady supply of reactants at low(er) supersaturation levels favoring the growth of well-developed euhedral faces detected in the *D. desulfuricans* cultures (Fig. 2J) (Raiswell, 1982). In contrast, the higher reactivity of ferrihydrite and ferric phosphate towards sulfide likely generated higher supersaturation levels leading to rapid crystal nucleation and growth of pyrite spherules, observed in many pure culture and abiotic incubations (Figs. 4J-K and N-O; 6K-O).

### 4.2 Bacteria in co-culture enhance or impair pyrite formation depending on the reactivity of the ferric compound present

Mackinawite was the dominant iron sulfide mineral in pre-diagenetic samples, consistent with its rapid formation from ferrous iron and sulfide (Duverger et al., 2020; Picard et al., 2018; Rickard & Luther, 2007) (Figs. 3A, 5A, 7A and 9A). Some of this mackinawite was oxidized, which can be attributed to the presence of ferric compounds and elemental S_(0)_/polysulfides (Bourdoiseau et al., 2008). Oxidized mackinawite is expected to undergo a solid-state transformation into greigite (Bourdoiseau et al., 2011; Csákberényi-Malasics et al., 2012). However, the unambiguous detection of greigite only in the *S. deleyianum* cultures corroborates previous findings that bacterial metabolisms can enhance iron sulfide transformations as opposed to purely abiotic incubations (Picard et al., 2018).

Pyrite precipitated under both biotic and abiotic conditions in the presence of ferric iron (goethite and ferric phosphate, Figs. S4 and 5A) and sulfide, consistent with previous observations (Duverger et al., 2020; Rickard, 1969). Given the apparent slow kinetics of the H_2_S-pathway at ambient temperatures (Picard et al., 2018), it is unlikely pyrite formed via this pathway. The surface of ferric oxides was shown to accelerate pyrite formation via the ferric-hydroxide-surface (FHS) pathway in conditions of high Fe and low S content (Wan et al., 2017). However, as the Fe:S ratio in our incubations was kept close to 1, we believe the FHS pathway might not have been the dominant mechanism of pyrite formation. Instead, since S_(0)_ was either externally added or formed through the reductive dissolution of ferric compounds, we hypothesize that the polysulfide pathway was driving pyrite precipitation prior to diagenesis (Duverger et al., 2020; Luther, 1991).

Contrary to our initial hypothesis, the actively metabolizing co-culture did not lead to a uniformly greater or faster pyrite production during the first weeks of the experiments (contrast between Fig. S4 and 5A). Instead, iron sulfide formation seemed to be modulated by the reactivity of the iron compounds, similar to the trend observed with mineral morphologies (see section 4.1). To initiate an internal cryptic S cycle (Fig. 1), biogenic sulfide produced by *D. desulfuricans* is expected to react with the ferric compound. In the case of less reactive goethite, its reductive dissolution by sulfide might proceed at a slower rate, resulting in the accumulation of free aqueous sulfide in the presence of Fe^2+^ and elemental S_(0)_ (Fig. 10A). Thus, elemental S_(0)_ could easily react with excess sulfide, giving rise to polysulfides which were hypothesized to be involved in pyrite formation in our incubations (see previous paragraph). Given that pyrite was readily detected in the co-cultures but not in pure cultures (Fig. S4), we theorize that the coupled metabolic activity of the co-culture created more favorable conditions for pyrite precipitation than the pure cultures alone. On the other hand, when the co-culture was grown with ferric phosphate, we expect that biogenic sulfide was immediately scavenged by this reactive ferric compound (Fig. 10B), resulting in rapid S_(0)_ and Fe^2+^ formation. However, further accumulation of elemental S_(0)_/polysulfides was likely prevented by its re-reduction to sulfide by *S. deleyianum*, creating a more reducing environment compared to the pure cultures. Such conditions would favor the preservation of metastable intermediates over pyrite, consistent with the detection of only mackinawite (co-culture in Fig. 5A). In contrast, elemental S_(0)_/polysulfides were likely present in higher amounts in pure cultures and abiotic incubations. In pure cultures of *D. desulfuricans*, S_(0)_ would have accumulated following the reductive dissolution of ferric phosphate, while *S. deleyianum* cultures were externally amended with it at the start of incubations (10 mM). These conditions likely resulted in a more oxidized environment as compared to co-cultures, which ultimately lead to pyrite formation (pure cultures and abiotic incubations in Fig. 5A). Overall, our results suggest that iron sulfide formation in our proposed cryptic S cycle is primarily governed by the reactivity of ferric minerals and bacterial metabolic rates. In other words, the presence of S-cycling bacteria alone is insufficient to explain the observed patterns in iron sulfide mineralogy. However, as we have not monitored the evolution of chemical species or pyritization rates over time, the mechanistic explanations laid out here remain to be verified by future investigations.

**Figure 10.**
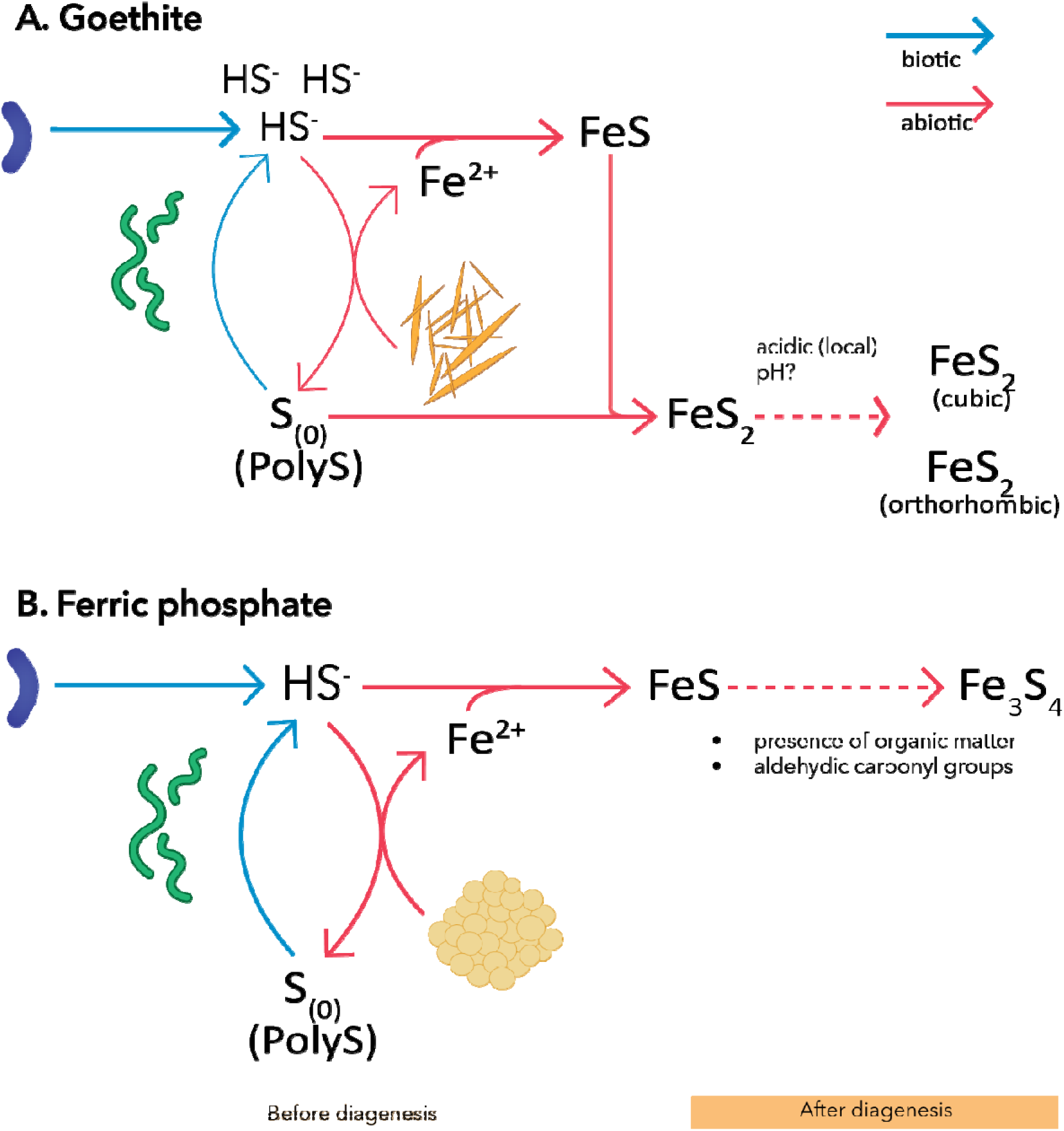
Schematic of the revised (hypothetical) internal cryptic S cycle mediated by the co-culture of *D. desulfuricans* and *S. deleyianum* in the presence of ferric compounds with different reactivity. For a more detailed mechanistic explanation, see the text in Section 4.2. The curved purple microbe is *D. desulfuricans*, while *S. deleyianum* is the green spirillum is. The thickness of arrows compares reaction rates (thicker arrows means faster reaction). Abbreviations: PolyS – polysulfides.

### 4.3 Co-occurrence of sulfate-and sulfur-reducing bacteria might promote the preservation of metastable iron sulfides during diagenesis

Simulated diagenesis chemically enhanced reaction rates in our abiotic or pure culture incubations. Heating experimentally added or endogenously produced elemental S_(0)_ and various metastable iron sulfides resulted in pyrite formation (goethite, ferric phosphate and ferrihydrite experiments, ferrous iron with S_(0)_) or further crystal growth where it was already present (goethite and ferric phosphate incubations). Given the presence of elemental S_(0)_, it is likely that pyrite continued to form via the polysulfide pathway as before diagenesis. In experiments with ferrous iron and no added S_(0)_, pyrite most probably precipitated via the H_2_S-pathway (Fe^2+^+*D. desulfuricans*, Fe^2+^+SO_4_^2-^+sulfide). Even though this pathway was thought to act slowly on laboratory timescales (Picard et al., 2018), pyrite formation here also appears to be accelerated by the increased temperature. Longer sulfidation experiments at ambient temperatures, as well as their comparison to heated samples, could verify this assumption.

In contrast, co-cultures were characterized by the persistence and, sometimes even (qualitative) dominance of metastable iron sulfides such as marcasite (goethite incubations; Figs. 2-3) and greigite (ferric phosphate, ferrihydrite and ferrous iron incubations; Figs. 4-9). Pyrite could not be unambiguously identified in all cases, suggesting that either it failed to form or its amount and/or crystallinity was too low to be detected by XRD or Raman spectroscopy. Since marcasite and greigite were not identified in any of the other corresponding pure culture or abiotic incubations prior to diagenesis, we hypothesize that the presence of co-cultured bacteria, as sources of organic matter, played a passive role in the appearance and persistence of these minerals. Pyrite was already present in goethite co-culture samples before diagenesis (Fig. S4). As marcasite is known to develop in acidic environments (Murowchick & Barnes, 1986), we hypothesize that locally acidic microniches may have acted as reactive sites for marcasite formation (co-culture in Fig. 3B, Fig 10A), similar as in pre-diagenetic samples (see section 4.1 above). For example, the thermal treatment of marine sediments at 85 °C, i.e. similar temperatures to our experiments, resulted in the increased abiotic production of acetic acid (Gan et al., 2025). It is therefore possible that the cell remains of co-cultures contained and/or released organic molecules that further decreased the (local) pH, leading to the simultaneous formation of post-diagenetic pyrite and marcasite. To verify this reaction mechanism, microscale multi-dimensional mapping techniques would be needed to track local pH changes over time (Hwang et al., 2016), which was not possible in our closed bottle incubations.

The post-diagenetic preservation of greigite can be explained by the delayed transformation kinetics of ferric compounds in the presence of microbial EPS (Wang et al., 2023) or organic matter (ThomasArrigo et al., 2020). Mackinawite, formed in our pre-diagenetic samples (co-cultures in 5A, 7A, 9A), might have thereby been prevented from transforming into pyrite, resulting in the accumulation of intermediate phases such as greigite (co-cultures in Figs. 5B, 7B and 9B; Fig 10B). Furthermore, aldehydic carbonyl groups can suppress pyrite formation in favor of greigite (Rickard et al., 2001). Even though we did not quantify these organic functional groups, their quantity might have been elevated in co-cultures due to the higher amount of starting biomass compared to pure cultures.

### 4.4 Abiotic disproportionation of elemental S_(0)_ produces spherulitic pyrite at elevated temperatures (80°C)

While microbes are thought to be the main producers of sulfide below 100 °C (Machel, 2001; van der Graaf et al., 2024), our results suggested abiotic sulfide formation from elemental S_(0)_ at 80 °C. We ruled out the contribution of sulfide from L-cysteine, a reactive thiol-containing compound, because no iron sulfides were detected in the controls amended with it (FP+*SO_4_^2^, Fh+*SO_4_^2-^) (Fig. 4B, 6B). Therefore, the most likely source of sulfide was either elemental S_(0)_ reduction or disproportionation. The stable concentration of formate over time excluded the possibility of S_(0)_ reduction (Fig. S10H-I), even though the redox potentials point toward the feasibility of this reaction under standard conditions (CO_2_/formate -431 mV, S/H_2_S -270 mV). Alternatively, elemental S_(0)_ might have been abiotically disproportionated according to the reaction stoichiometry performed by some bacteria (Reaction 1; Wu et al., 2025), which is supported by the gradual increase in thiosulfate, but not sulfate concentrations over time (up to 1 mM, Fig. S10I). In our system, sulfide would have been continuously removed by the ferric iron minerals keeping the disproportionation reaction thermodynamically favorable (Thamdrup et al., 1993). Taken together, these observations indicate that the abiotic disproportionation of S_(0)_ can produce sulfide, and subsequently, spherulitic pyrite at temperatures present in diagenetic or hydrothermal habitats (Fig. S10).

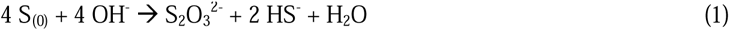

It is well-known that microbes are capable of disproportionating elemental S_(0)_; with an optimum growth temperature often ∼ 30 °C (Finster et al., 1998; Thamdrup et al., 1993) but reaching up to ∼ 74 °C in the case of (hyper)thermophilic bacteria (Slobodkin et al., 2012). In contrast, elemental S_(0)_ has been shown to be chemically disproportionated at elevated temperatures (≥ 200 °C) (Maffeis et al., 2024; Robinson, 1973; Yuan et al., 2013). To our knowledge, abiotic S_(0)_ disproportionation was not previously experimentally verified at temperatures as low as 80 °C. In earlier experiments, it was postulated that this reaction could explain the increase in sulfide concentrations at similar temperatures, even though sulfate/thiosulfate concentrations were not measured (Belkin et al., 1985). Overall, our results extend the feasibility of abiotic iron sulfide formation to temperature regimes where sulfide is usually thought to have a microbial origin (van der Graaf et al., 2024).

### 4.5 Abiotic sulfide production at elevated (80 °C) temperatures in the presence of ferrous iron

Abiotic or thermochemical reduction of sulfate is believed to become feasible at high temperatures (starting from 100-140 °C) and become a geologically important process at ≥ 200 °C (Machel, 2001; Ohmoto & Lasaga, 1982). The surprising development of iron sulfides, especially pyrite, in our controls with only Fe^2+^ and SO_4_^2-^at 80 °C was therefore further investigated. While these initial results were reproducible, iron sulfides also developed in sulfate-free incubations with only ferrous iron. The medium did not contain any L-cysteine or elemental S_(0)_ (see above) which could have either released sulfide or be disproportionated over time. However, we used an organosulfonate MOPS (∼ 15 mM) buffer in all of our incubations to maintain a pH of ∼ 7.2. While MOPS is regarded as a Good’s buffer, thus expected to be inert, there is insufficient information about its stability and potential abiotic transformations at 80 °C over long time periods.

Alternatively, butyl rubber stoppers commonly used in anaerobic microbiology can contain sulfur after the vulcanization process. While this cross-linked sulfur can be leached by aggressive methods (Zheng et al., 2020), our controls (FP+SO_4_^2^, Fh+SO_4_^2-^, FP+*SO_4_^2^, Fh+*SO_4_^2-^) supplemented with various ferric iron compounds remained visually and mineralogically unchanged after simulated diagenesis (Figs. 4P, 5B, 6P, 7B), suggesting negligible sulfide release from the stoppers. Thus, we posit that this process was of diminutive importance in our overall experimental setup.

In addition to iron sulfides, we could also detect magnetite in our controls. Magnetite was previously shown to form via the Schikorr reaction (Truong et al., 2025), as shown in Reaction 2.

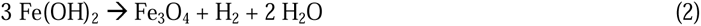

The Schikorr reaction is thought to be feasible at low temperatures (< 100 °C), potentially resulting in the concomitant release of H_2_ gas. Although H_2_ was not measured, it might have been involved in abiotic sulfide production. Furthermore, as magnetite is an electrically conductive mineral (Radoń et al., 2018), it may have facilitated the oxidation of H_2_ during this process (van der Graaf et al., 2024).

Overall, organosulfonate (and potentially sulfate, which cannot be excluded by our current analyses) transformations are the most likely source of abiotic sulfide formation in the presence of ferrous iron at 80 °C. Since these compounds are of environmental relevance in hydrothermal systems (Rose et al., 2001; Sajkowski et al., 2022) or potential substrates sustaining heterotrophic communities (Chen et al., 2022), this phenomenon warrants further study. Future experiments should therefore focus on deciphering the reaction mechanism as a function of temperature and measuring H_2_ gas that would provide support for the Schikorr reaction.

### 4.6 Implications for natural environments

Our results demonstrate that ferric mineral reactivity exerts a significant control on pyrite morphology during diagenesis, most probably through its influence on pyrite supersaturation. We add to a growing body of evidence indicating that pyrite morphology reflects the physicochemical conditions under which pyritization occurs rather than biological factors. Previous studies have shown that elemental S_(0)_ (Runge et al., 2024), as well as high Eh and/or low pH (Butler & Rickard, 2000) might increase pyrite supersaturation levels, resulting thus in the rapid formation of pyrite spherules or framboids. We have not explicitly measured pyritization rates as a function of ferric compound reactivity or presence of S_(0)_. However, it is likely that pyrite formed at a faster rate in our incubations amended with elemental S_(0)_ through the formation of polysulfides, as was shown in previous studies (Runge et al., 2024). The presence of organic matter was hypothesized to impede pyritization by complexing reactive intermediates such as S_(0)_ and FeS (Cosmidis et al., 2019; Picard et al., 2019). However, our abiotic and pure culture incubations contained similar crystal habits, suggesting a stronger control of ferric iron reactivity and diagenetic heating over the presence of bacterial cell remains under the conditions tested. Taken together, we expect that ferric iron reactivity, Eh/pH, the presence of S_(0)_ and organic matter may act collectively in regulating supersaturation levels in natural habitats, thereby influencing both pyritization rates and morphology.

Our results further highlight that iron sulfides preserved in sediments may reflect local interactions between diagenetic processes and microbial communities (Pasquier et al., 2025). Given the formation of pyrite in both pure culture and abiotic incubations, we propose that pyritization is a primarily chemically controlled process that is enhanced by temperature during sediment diagenesis. Even though microbes are the primary source of sulfide in Earth surface environments, their further presence might not be mechanistically necessary for the development of pyrite. However, by serving as an organic matter source during diagenesis they likely continue to influence this process in natural habitats, as observed with the appearance of marcasite and greigite in our post-diagenetic co-culture incubations. The occurrence of marcasite and greigite might have important implications for paleoenvironmental reconstructions. Marcasite is considered less stable than pyrite, yet it becomes the dominant iron disulfide mineral in acidic habitats and is therefore commonly used as a pH paleoindicator (Lounejeva et al., 2021). Greigite is a ferrimagnetic intermediate in the polysulfide pathway (Hunger & Benning, 2007; Roberts et al., 2011) that is generally considered unlikely to be preserved in the rock record (Berner, 1983). Nevertheless, experimental and thermodynamic studies have shown that decreased sulfate and organic matter content, together with an increase in reactive iron can promote its preservation (Blanchet et al., 2009; Fu et al., 2015; Kao et al., 2004; Rickard et al., 2024). Accordingly, the appearance of greigite in sedimentary layers is often used as a paleoindicator to track sea level and salinity changes which are sensitive to fluctuations in sulfate or reactive iron content (Blanchet et al., 2009; Chen et al., 2021; Liu et al., 2017). However, our results indicate that greigite and marcasite can continue to precipitate/grow in deeper sedimentary layers in the presence of complex microbial S-cycling communities. As a result, the primary mineralogical signatures formed at the time of the sediment deposition might be modified or overprinted during burial diagenesis (Ron et al., 2007), thereby complicating reconstructions of past environments. To further expand on the diagenetic formation and persistence of greigite and marcasite, future studies should consider exposing different types of organic matter (e.g. plant material, complex microbial communities) to experimental diagenetic conditions.

It has long been believed that a firm temperature gap exists between microbially and abiotically produced sulfide. However, we experimentally demonstrated the precipitation of iron sulfides in a previously overlooked abiotic reaction occurring at elevated temperatures (80 °C): disproportionation of S_(0)_. Elemental S_(0)_ is speculated to have been widely present on ancient Earth due to the photolysis of SO_2_ (Catling & Zahnle, 2020; Dewitt et al., 2010; Ono, 2026). On modern Earth, S_(0)_ is present wherever (a)biotic recycling of S compounds occurs (Dahl, 2020). Given that temperatures of up to 80 °C are common near hydrothermal vents and in deeper sediment layers, the abiotic formation of iron sulfides via the mentioned reaction is thus feasible in such conditions. Consequently, the contribution of abiotically produced sulfide in iron sulfide precipitation might be more significant and widespread than previously assumed. Future studies should therefore aim to disentangle the relative contribution of microbially and abiotically made sulfide in these intermediate temperature zones, especially with regard to biosignatures. Since the microbial reduction of sulfate is known to create large isotope fractionations (Sim et al., 2023), measuring the S isotope effect associated with the abiotic disproportionation of S_(0)_ may offer a useful geochemical tool for differentiating between abiotically and microbially generated iron sulfides.

Finally, our findings question the use of spherulitic pyrite as a robust biosignature. Spherulitic pyrite was previously interpreted as evidence for microbial activity because its formation was thought to require complex organic matter derived from microbial cell remains (Duverger et al., 2021; Truong et al., 2024). In contrast, our abiotic incubations contained only simple organic acids such as lactate, formate and acetate that served as electron donors/nutrients in the bacterial incubations. Therefore, we speculate that the presence of spherulitic pyrite in the rock record might not be indicative of microbial presence but rather reflects on environmental conditions with the reactivity of ferric compounds highlighted in our work. On the other hand, in some co-culture samples (ferric phosphate and goethite) the appearance of greigite and marcasite resulted in the development of less distinguishable (non-spherulitic) crystal habits. Consequently, defining morphological criteria for the biogenicity of iron sulfides is likely to be challenging due to several factors: (1) abiotic pyrite having well-defined morphologies and (2) mixed microbial communities causing a change in iron sulfide mineralogy and so crystal shape. Taken together, these findings suggest that pyrite morphology provides an insufficient basis for assessing biogenicity and should be interpreted alongside mineralogical, isotopic, and organic geochemical criteria (Picard et al., 2021; Truong et al., 2023) within a multiproxy framework, as suggested previously.

## 5 Conclusions

Our study demonstrates that iron sulfide evolution is governed by interactions between microbial S cycling, iron mineral reactivity and diagenetic transformations. Contrary to our initial hypothesis, the S-cycling co-culture did not universally promote pyritization but instead favored the preservation of metastable minerals such as greigite and marcasite after diagenesis. Thus, complex microbial communities subjected to burial diagenesis in natural sediments might confound our paleo(magnetic) reconstructions and overall understanding of the evolution of the S cycle. We also highlighted that the boundary between microbially and abiotically generated sulfide may be less distinct than previously assumed in sedimentary and hydrothermal habitats at relatively low temperatures (80°C). Lastly, we showed that the morphology of diagenetic pyrite depended on the reactivity of iron minerals rather than the presence of bacteria, thus challenging assumptions that pyrite morphology can serve as an indicator for microbial presence. Consequently, to reconstruct paleoenvironments and define candidate biosignatures based on iron sulfides, future studies should explicitly consider the potential of both diagenetic alternation and abiotic S transformations.

## Notes

https://zenodo.org/records/20797468

### Competing Interest Statement

The authors have declared no competing interest.

### Funding

- Swiss National Science Foundation: 212159

## References

Aller, R. (2014). Sedimentary Diagenesis, Depositional Environments, and Benthic Fluxes. In H. D. Holland & K. K. Turekian (Eds.), Treatise on Geochemistry (Second Edition, Vol. 8, pp. 293–334). Elsevier.

Avetisyan, K., Zweig, I., Luther, G. W., & Kamyshny, A. (2021). Kinetics and mechanism of polysulfides and elemental sulfur formation by a reaction between hydrogen sulfide and δ-MnO2. Geochimica et Cosmochimica Acta, 313, 21–37. 10.1016/j.gca.2021.08.022

Belkin, S., Wirsen, C. O., & Jannasch, H. W. (1985). Biological and Abiological Sulfur Reduction at High Temperatures. Applied and Environmental Microbiology, (49), 1057–1061. 10.1128/aem.49.5.1057-1061.1985

Benning, L. G., Wilkin, R. T., & Konhauser, K. O. (1999). Iron monosulphide stability: Experiments with sulphate reducing bacteria. In H. Armannsson (Ed.), Geochemistry of the Earth’s Surface (1st ed., pp. 429–432). CRC Press. 10.1201/9781003761938

Berg, J. S., Duverger, A., Cordier, L., Laberty-Robert, C., Guyot, F., & Miot, J. (2020). Rapid pyritization in the presence of a sulfur/sulfate-reducing bacterial consortium. Scientific Reports, 10(1), 8264. 10.1038/s41598-020-64990-6

Berner, R. A. (1983). Sedimentary pyrite formation: An update*. Geochimica et Cosmochimica Acta, 48, 605– 615. 10.1016/0016-7037(84)90089-9

Blanchet, C. L., Thouveny, N., & Vidal, L. (2009). Formation and preservation of greigite (Fe3S4) in sediments from the Santa Barbara Basin: Implications for paleoenvironmental changes during the past 35 ka. Paleoceanography, 24(2), PA2224. 10.1029/2008PA001719

Böttcher, M. E., Smock, A. M., & Cypionka, H. (1998). Sulfur isotope fractionation during experimental precipitation of iron (II) and manganese (II) sulfide at room temperature. Chemical Geology, 146, 127–134. 10.1016/S0009-2541(98)00004-7

Bourdoiseau, J. A., Jeannin, M., Rémazeilles, C., Sabot, R., & Refait, P. (2011). The transformation of mackinawite into greigite studied by Raman spectroscopy. Journal of Raman Spectroscopy, 42(3), 496–504. 10.1002/jrs.2729

Bourdoiseau, J. A., Jeannin, M., Sabot, R., Rémazeilles, C., & Refait, P. (2008). Characterisation of mackinawite by Raman spectroscopy: Effects of crystallisation, drying and oxidation. Corrosion Science, 50(11), 3247– 3255. 10.1016/j.corsci.2008.08.041

Butler, I. B., & Rickard, D. (2000). Framboidal pyrite formation via the oxidation of iron (II) monosulfide by hydrogen sulphide. Geochimica et Cosmochimica Acta, 64(15), 2665–2672.

Canfield, D. E. (1989). Reactive iron in marine sediments. Geochimica et Cosmochimica Acta, 53, 619–632. 10.1016/0016-7037(89)90005-7

Canfield, D. E., Thamdrup, B., & Fleischer, S. (1998). Isotope fractionation and sulfur metabolism by pure and enrichment cultures of elemental sulfur-disproportionating bacteria. Limnology and Oceanography, 43(2), 253–264. 10.4319/lo.1998.43.2.0253

Catling, D. C., Krissansen-Totton, J., Kiang, N. Y., Crisp, D., Robinson, T. D., DasSarma, S., Rushby, A. J., Del Genio, A., Bains, W., & Domagal-Goldman, S. (2018). Exoplanet Biosignatures: A Framework for Their Assessment. Astrobiology, 18(6), 709–738. 10.1089/ast.2017.1737

Catling, D. C., & Zahnle, K. J. (2020). The Archean atmosphere. Science Advances, 6, eaax1420. https://www.science.org

Ceriotti, G., Bosco-Santos, A., & Berg, J. S. (2025). Oxic microbial ferrihydrite reduction rates of Shewanella oneidensis and the potential for Fe mobilization in oxic sediments. Scientific Reports, 15, 31343. 10.1038/s41598-025-16963-w

Chen, K. Y., & Gupta, S. K. (1973). Formation of polysulfides in aqueous solutions. Environmental Letters, 4(3), 187–200.

Chen, X., Han, Y., Chen, Q., Lin, H., Lin, S., Wang, D., & Tang, K. (2022). Ubiquitous Occurrence of a Biogenic Sulfonate in Marine Environment. Sustainability, 14(3), 1240. 10.3390/su14031240

Chen, Y., Zhang, W., Nian, X., Sun, Q., Ge, C., Hutchinson, S. M., Cheng, Q., Wang, F., Chen, J., & Zhao, X. (2021). Greigite as an Indicator for Salinity and Sedimentation Rate Change: Evidence From the Yangtze River Delta, China. Journal of Geophysical Research: Solid Earth, 126(3), e2020JB021085. 10.1029/2020JB021085

Chi Fru, E., Callac, N., Posth, N. R., Argyraki, A., Ling, Y. C., Ivarsson, M., Broman, C., & Kilias, S. P. (2018). Arsenic and high affinity phosphate uptake gene distribution in shallow submarine hydrothermal sediments. Biogeochemistry, 141(1), 41–62. 10.1007/s10533-018-0500-8

Cornell, R. M., & Schwertmann, U. (2003). The Iron Oxides: Structure, Properties, Reactions, Occurences and Uses. Wiley-VCH. 10.1002/3527602097

Cosmidis, J., Nims, C. W., Diercks, D., & Templeton, A. S. (2019). Formation and stabilization of elemental sulfur through organomineralization. Geochimica et Cosmochimica Acta, 247, 59–82. 10.1016/j.gca.2018.12.025

Csákberényi-Malasics, D., Rodriguez-Blanco, J. D., Kis, V. K., Rečnik, A., Benning, L. G., & Pósfai, M. (2012). Structural properties and transformations of precipitated FeS. Chemical Geology, 294-295, 249–258. 10.1016/j.chemgeo.2011.12.009

Dahl, C. (2020). A biochemical view on the biological sulfur cycle. In L. N. L. Piet (Ed.), Environmental Technologies to Treat Sulfur Pollution: Principles and Engineering (2nd edition, pp. 55–96). IWA Publishing. 10.2166/9781789060966_0055

Dewitt, H. L., Hasenkopf, C. A., Trainer, M. G., Farmer, D. K., Jimenez, J. L., McKay, C. P., Toon, O. B., & Tolbert, M. A. (2010). The formation of sulfate and elemental sulfur aerosols under varying laboratory conditions: Implications for early Earth. Astrobiology, 10(8), 773–781. 10.1089/ast.2009.9455

Duverger, A., Berg, J. S., Busigny, V., Guyot, F., Bernard, S., & Miot, J. (2020). Mechanisms of Pyrite Formation Promoted by Sulfate-Reducing Bacteria in Pure Culture. Frontiers in Earth Science, 8, 588310. 10.3389/feart.2020.588310

Duverger, A., Bernard, S., Viennet, J. C., Miot, J., & Busigny, V. (2021). Formation of Pyrite Spherules From Mixtures of Biogenic FeS and Organic Compounds During Experimental Diagenesis. Geochemistry, Geophysics, Geosystems, 22(10). 10.1029/2021GC010056

Dzieniszewska, A., Kyziol-Komosinska, J., & Pająk, M. (2020). Adsorption and bonding strength of chromium species by ferrihydrite from acidic aqueous solutions. PeerJ, 2020(8), e9324. 10.7717/peerj.9324

Eder, S. H. K., Gigler, A. M., Hanzlik, M., & Winklhofer, M. (2014). Sub-micrometer-scale mapping of magnetite crystals and sulfur globules in magnetotactic bacteria using confocal raman micro-spectrometry. PLoS ONE, 9(9), e107356. 10.1371/journal.pone.0107356

Findlay, A. J., & Kamyshny, A. (2017). Turnover rates of intermediate sulfur species (Sx2-, S0, S2O32-, S4O62-, SO32-) in anoxic freshwater and sediments. Frontiers in Microbiology, 8, 2551. 10.3389/fmicb.2017.02551

Finster, K., Liesack, W., & Thamdrup, B. (1998). Elemental Sulfur and Thiosulfate Disproportionation by Desulfocapsa sulfoexigens sp. nov., a New Anaerobic Bacterium Isolated from Marine Surface Sediment. Applied and Environmental Microbiology, 64(1), 119–125. 10.1128/AEM.64.1.119-125.1998

Fortin, D., Southam, G., & Beveridge, T. J. (1994). Nickel sulfide, iron-nickel sulfide and iron sulfide precipitation by a newly isolated Desulfotomaculum species and its relation to nickel resistance. FEMS Microbiology Ecology, 14, 121–132. 10.1111/j.1574-6941.1994.tb00099.x

Fu, C., Bloemendal, J., Qiang, X., Hill, M. J., & An, Z. (2015). Occurrence of greigite in the Pliocene sediments of Lake Qinghai, China, and its paleoenvironmental and paleomagnetic implications. Geochemistry, Geophysics, Geosystems, 16(5), 1293–1306. 10.1002/2014GC005677

Gan, S., Heuer, V. B., Schmidt, F., Wörmer, L., Wang, F., Adhikari, R. R., Hatcher, P., Pearson, A., & Hinrichs, K.-U. (2025). Moderate heating renders 7.8-million-year-old sedimentary organic matter bioavailable. Science Advances, 11(34), eadw8638.

Gramp, J. P., Bigham, J. M., Jones, F. S., & Tuovinen, O. H. (2010). Formation of Fe-sulfides in cultures of sulfate-reducing bacteria. Journal of Hazardous Materials, 175(1–3), 1062–1067. 10.1016/j.jhazmat.2009.10.119

Halevy, I., Peters, S. E., & Fischer, W. W. (2012). Sulfate Burial Constraints on the Phanerozoic Sulfur Cycle. Science, 337(6092), 331–334. 10.1126/science.1219703

Hansel, C. M., Lentini, C. J., Tang, Y., Johnston, D. T., Wankel, S. D., & Jardine, P. M. (2015). Dominance of sulfur-fueled iron oxide reduction in low-sulfate freshwater sediments. ISME Journal, 9(11), 2400–2412. 10.1038/ismej.2015.50

Herbert, R. B. J., Benner, S. G., Pratt, A. R., & Blowes, D. W. (1998). Surface chemistry and morphology of poorly crystalline iron sulfides precipitated in media containing sulfate-reducing bacteria. Chemical Geology, 144, 87–97. 10.1016/S0009-2541(97)00122-8

Hidalgo, G., Burns, A., Herz, E., Hay, A. G., Houston, P. L., Wiesner, U., & Lion, L. W. (2009). Functional tomographic fluorescence imaging of pH microenvironments in microbial biofilms by use of silica nanoparticle sensors. Applied and Environmental Microbiology, 75(23), 7426–7435. 10.1128/AEM.01220-09

Hoffmann, T. D., Reeksting, B. J., & Gebhard, S. (2021). Bacteria-induced mineral precipitation: A mechanistic review. Microbiology, 167(4), 001049. 10.1099/mic.0.001049

Hunger, S., & Benning, L. G. (2007). Greigite: A true intermediate on the polysulfide pathway to pyrite. Geochemical Transactions, 8. 10.1186/1467-4866-8-1

Hwang, G., Liu, Y., Kim, D., Sun, V., Aviles-Reyes, A., Kajfasz, J. K., Lemos, J. A., & Koo, H. (2016). Simultaneous spatiotemporal mapping of in situ pH and bacterial activity within an intact 3D microcolony structure. Scientific Reports, 6, 32841. 10.1038/srep32841

Kao, S.-J., Horng, C.-S., Roberts, A. P., & Liu, K.-K. (2004). Carbon-sulfur-iron relationships in sedimentary rocks from southwestern Taiwan: Influence of geochemical environment on greigite and pyrrhotite formation. Chemical Geology, 203(1–2), 153–168. 10.1016/j.chemgeo.2003.09.007

Koblitz, J., Halama, P., Spring, S., Thiel, V., Baschien, C., Hahnke, R. L., Pester, M., Overmann, J., & Reimer, L. C. (2023). MediaDive: the expert-curated cultivation media database. Nucleic Acids Research, 51(1), D1531–D1538. 10.1093/nar/gkac803

Konhauser, K. O. (1998). Diversity of bacterial iron mineralization. Earth-Science Reviews, 43, 91–121. 10.1016/S0012-8252(97)00036-6

Kubeneck, L. J., Fantappiè, G., Notini, L., Rothwell, K. A., ThomasArrigo, L. K., & Kretzschmar, R. (2025). Interplay of Fe and S biogeochemistry shapes in situ iron mineral transformations in contrasting intertidal sediments. Environmental Science: Processes and Impacts, 27(12), 3903–3916. 10.1039/d5em00213c

Lentini, C. J., Wankel, S. D., & Hansel, C. M. (2012). Enriched iron(III)-reducing bacterial communities are shaped by carbon substrate and iron oxide mineralogy. Frontiers in Microbiology, 3, 404. 10.3389/fmicb.2012.00404

Li, Y.-L., Vali, H., Yang, J., Phelps, T. J., & Zhang, C. L. (2006). Reduction of iron oxides enhanced by a sulfate-reducing bacterium and biogenic H2S. Geomicrobiology Journal, 23(2), 103–117. 10.1080/01490450500533965

Liu, S., Krijgsman, W., Dekkers, M. J., & Palcu, D. (2017). Early diagenetic greigite as an indicator of paleosalinity changes in the middle Miocene Paratethys Sea of central Europe. *Geochemistry, Geophysics*, Geosystems, 18(7), 2634–2645. 10.1002/2017GC006988

Liu, Y., Li, Z., You, Y., Zheng, X., & Wen, J. (2017). Synthesis of different structured FePO4 for the enhanced conversion of methyl cellulose to 5-hydroxymethylfurfural. RSC Advances, 7(81), 51281–51289. 10.1039/c7ra09186a

Lounejeva, E., Steadman, J. A., Rodemann, T., Large, R. R., Danyushevsky, L., Mantle, D., Grice, K., & Algeo, T. J. (2021). Marcasite at the Permian-Triassic Transition: A Potential Indicator of Hydrosphere Acidification. In R. E. Ernst, A. J. Dickson, & A. Bekker (Eds.), Large Igneous Provinces: A Driver of Global Environmental and Biotic Changes (1st ed., pp. 377–399). American Geophysical Union and John Wiley and Sons, Inc. 10.1002/9781119507444.ch16

Luther, G. W. I. (1991). Pyrite synthesis via polysulfide compounds. Geochimica et Cosmochimica Acta, 55, 2839–2849. 10.1016/0016-7037(91)90449-F

Machel, H. G. (2001). Bacterial and thermochemical sulfate reduction in diagenetic settings - old and new insights. Sedimentary Geology, 140(1–2), 143–175. 10.1016/S0037-0738(00)00176-7

Maffeis, A., Frezzotti, M. L., Connolly, J. A. D., Castelli, D., & Ferrando, S. (2024). Sulfur disproportionation in deep COHS slab fluids drives mantle wedge oxidation. Science Advances, 10(12), eadj2770. 10.1126/sciadv.adj2770

Mansor, M., Duverger, A., Pasquier, V., Gorlas, A., Guyot, F., Berg, J. S., Marin-Carbonne, J., Cosmidis, J., & Picard, A. (2025). Biogenic pyrite and metastable iron sulfides: Emerging formation pathways and geological and societal relevance. Geo-Bio Interfaces, 2, e6. 10.1180/gbi.2024.9

Marin-Carbonne, J., Decraene, M.-N., Dupeyron, J., Alleon, J., Pasquier, V., Remusat, L., Thomazo, C., Olivier, N., Benzerara, K., & Bernard, S. (2023). Micropyrite: a promising biosignature? Insights from modern and ancient sediments. Goldschmidt Conference.

Mirabello, G., GoodSmith, M., Bomans, P. H. H., Stegbauer, L., Joester, D., & de With, G. (2021). Iron phosphate mediated magnetite synthesis: a bioinspired approach. Chemical Science, 12(27), 9458–9465. 10.1039/d0sc07079c

Mirvaux, B., Recham, N., Miot, J., Courty, M., Bernard, S., Beyssac, O., Davoisne, C., Sougrati, M., Demortière, A., Guyot, F., Tarascon, J. M., & Larcher, D. (2016). Iron Phosphate/Bacteria Composites as Precursors for Textured Electrode Materials with Enhanced Electrochemical Properties. Journal of The Electrochemical Society, 163(10), A2139–A2148. 10.1149/2.0101610jes

Murowchick, J. B., & Barnes, H. L. (1986). Marcasite precipitation from hydrothermal solutions. Geochimica et Cosmochimica Acta, 50(12), 2615–2629. 10.1016/0016-7037(86)90214-0

Murowchick, J. B., & Barnes, H. Lloyd. (1987). Effects of temperature and degree of supersaturation on pyrite morphology. American Mineralogist, 72, 1241–1250.

Nabeh, N., Brokaw, C., & Picard, A. (2022). Quantification of Organic Carbon Sequestered by Biogenic Iron Sulfide Minerals in Long-Term Anoxic Laboratory Incubations. Frontiers in Microbiology, 13, 662219. 10.3389/fmicb.2022.662219

Nguyen, A. T., Goswami, S., Ferracane, J., & Koley, D. (2022). Real-time monitoring of the pH microenvironment at the interface of multispecies biofilm and dental composites. Analytica Chimica Acta, 1201, 339589. 10.1016/j.aca.2022.339589

Ohmoto, H., & Lasaga, A. C. (1982). Kinetics of reactions between aqueous sulfates and sulfides in hydrothermal systems. Geochimica et Cosmochimica Acta, 46(10), 1727–1745.

Ono, S. (2026). Photochemistry of Sulfur Dioxide and the Origin of Mass-Independent Isotope Fractionation in Earth’s Atmosphere. The Annual Review of Earth and Planetary Sciences Is Online at Earth.Annualreviews.Org Downloaded from Www.Annualreviews.Org. Guest, 45, 301–329. 10.1146/annurev-earth-060115

Pasquier, V., Marin-Carbonne, J., Giunta, T., Ruffine, L., & Halevy, I. (2025). Microscale iron and sulphur isotopic compositions reveal pyritization pathways during early diagenesis. Communications Earth and Environment, 6(1), 248. 10.1038/s43247-025-02213-4

Peiffer, S., Behrends, T., Hellige, K., Larese-Casanova, P., Wan, M., & Pollok, K. (2015). Pyrite formation and mineral transformation pathways upon sulfidation of ferric hydroxides depend on mineral type and sulfide concentration. Chemical Geology, 400, 44–55. 10.1016/j.chemgeo.2015.01.023

Peña, F. de la, Prestat, E., Fauske, V. T., Burdet, P., Furnival, T., Jokubauskas, P., Nord, M., Ostasevicius, T., MacArthur, K. E., Johnstone, D., Sarahan, M., Lähnemann, J., Taillon, J., pquinn-dls, Aarholt, T., Migunov, V., Eljarrat, A., Caron, J., Mazzucco, S., … Myers, J. C. (2020). hyperspy/hyperspy: Release v1.6.1.

Picard, A., Gartman, A., Clarke, D. R., & Girguis, P. R. (2018). Sulfate-reducing bacteria influence the nucleation and growth of mackinawite and greigite. Geochimica et Cosmochimica Acta, 220, 367–384. 10.1016/j.gca.2017.10.006

Picard, A., Gartman, A., Cosmidis, J., Obst, M., Vidoudez, C., Clarke, D. R., & Girguis, P. R. (2019). Authigenic metastable iron sulfide minerals preserve microbial organic carbon in anoxic environments. Chemical Geology, 530, 119343. 10.1016/j.chemgeo.2019.119343

Picard, A., Gartman, A., & Girguis, P. R. (2016). What do we really know about the role of microorganisms in iron sulfide mineral formation? Frontiers in Earth Science, 4, 68. 10.3389/feart.2016.00068

Picard, A., Gartman, A., & Girguis, P. R. (2021). Interactions between Iron Sulfide Minerals and Organic Carbon: Implications for Biosignature Preservation and Detection. Astrobiology, 21(5), 587–604. 10.1089/ast.2020.2276

Raiswell, R. (1982). Pyrite texture, isotopic composition and the availability of iron. American Journal of Science, 282, 1244–1263.

Rickard, D. (1969). The microbiological formation of iron sulfides. Stockholm Contributions in Geology, 20(3), 49–66. https://www.researchgate.net/publication/284577911

Rickard, D. (1997). Kinetics of pyrite formation by the H2S oxidation of iron (II) monosulfide in aqueous solutions between 25 and 125°C: The rate equation. Geochimica et Cosmochimica Acta, 61(1), 115–134.

Rickard, D. (2012). Sedimentary Pyrite. In Sulfidic Sediments and Sedimentary Rocks (Vol. 65, pp. 233–285). Elsevier. 10.1016/B978-0-444-52989-3.00006-4

Rickard, D., Butler, I. B., & Oldroyd, A. (2001). A novel iron sulphide mineral switch and its implications for Earth and planetary science. Earth and Planetary Science Letters, 189, 85–91. 10.1016/S0012-821X(01)00352-1

Rickard, D., & Luther, G. W. (1997). Kinetics of pyrite formation by the H2S oxidation of iron (II) monosulfide in aqueous solutions between 25 and 125°C: The mechanism. Geochimica et Cosmochimica Acta, 61(1), 135–147.

Rickard, D., & Luther, G. W. I. (2007). Chemistry of iron sulfides. In Chemical Reviews (Vol. 107, Number 2, pp. 514–562). 10.1021/cr0503658

Rickard, D., Roberts, A. P., & Navrotsky, A. (2024). Sedimentary Greigite Formation. American Journal of Science, 324. 10.2475/001c.121855

Rickard, D. T. (1974). Kinetics and mechanisms of the sulfidation of goethite. American Journal of Science, 274, 941–952.

Roberts, A. P., Chang, L., Rowan, C. J., Horng, C. S., & Florindo, F. (2011). Magnetic properties of sedimentary greigite (Fe3S4): An update. Reviews of Geophysics, 49(1), RG1002. 10.1029/2010RG000336

Robinson, B. W. (1973). Sulphur isotope equilibrium during sulphur hydrolysis at high temperatures. Earth and Planetary Science Letters, 18, 443–450.

Ron, H., Nowaczyk, N. R., Frank, U., Schwab, M. J., Naumann, R., Striewski, B., & Agnon, A. (2007). Greigite detected as dominating remanence carrier in Late Pleistocene sediments, Lisan formation, from Lake Kinneret (Sea of Galilee), Israel. Geophysical Journal International, 170(1), 117–131. 10.1111/j.1365-246X.2007.03425.x

Rose, P. E., Benoit, W. R., & Kilbourn, P. M. (2001). The application of the polyaromatic sulfonates as tracers in geothermal reservoirs. Geothermics, 30, 617–640. www.elsevier.com/locate/geothermics

Runge, E. A., Mansor, M., Kappler, A., & Duda, J. P. (2023). Microbial biosignatures in ancient deep-sea hydrothermal sulfides. Geobiology, 21(3), 355–377. 10.1111/gbi.12539

Runge, E., Mansor, M., Chiu, T. H., Shuster, J., Fischer, S., Kappler, A., & Duda, J. P. (2024). Hydrothermal sulfidation of biogenic magnetite produces framboid-like pyrite. Communications Earth and Environment, 5(1), 252. 10.1038/s43247-024-01400-z

Sajkowski, L., Seward, T. M., Mountain, B. W., & Marynowski, L. (2022). The stability of polyaromatic naphthalene sulfonates in hydrothermal solutions to 330 °C at equilibrium saturated vapour pressure. Geothermics, 104, 102437. 10.1016/j.geothermics.2022.102437

Schoonen, M. A. A. (2004). Mechanisms of sedimentary pyrite formation. In J. P. Amend, K. J. Edwards, & T. W. Lyons (Eds.), Sulfur Biogeochemistry - Past and Present (Vol. 379, pp. 117–134). Geological Society of America.

Sekerci, F., Fischer, S., Joshi, P., Peiffer, S., Kappler, A., & Mansor, M. (2025). Sulfur microenvironments as hotspots for biogenic pyrite formation. Scientific Reports, 15(1), 20148. 10.1038/s41598-025-05178-8

Sim, M. S., Woo, D. K., Kim, B., Jeong, H., Joo, Y. J., Hong, Y. W., & Choi, J. Y. (2023). What Controls the Sulfur Isotope Fractionation during Dissimilatory Sulfate Reduction? In ACS Environmental Au (Vol. 3, Number 2, pp. 76–86). American Chemical Society. 10.1021/acsenvironau.2c00059

Slobodkin, A. I., Reysenbach, A. L., Slobodkina, G. B., Baslerov, R. V., Kostrikina, N. A., Wagner, I. D., & Bonch-Osmolovskaya, E. A. (2012). Thermosulfurimonas dismutans gen. nov., sp. nov., an extremely thermophilic sulfur-disproportionating bacterium from a deep-sea hydrothermal vent. International Journal of Systematic and Evolutionary Microbiology, 62(11), 2565–2571. 10.1099/ijs.0.034397-0

Straub, K. L., & Schink, B. (2004). Ferrihydrite-dependent growth of Sulfurospirillum deleyianum through electron transfer via sulfur cycling. Applied and Environmental Microbiology, 70(10), 5744–5749. 10.1128/AEM.70.10.5744-5749.2004

Sun, J., Mailloux, B. J., Chillrud, S. N., van Geen, A., Thompson, A., & Bostick, B. C. (2018). Simultaneously quantifying ferrihydrite and goethite in natural sediments using the method of standard additions with X-ray absorption spectroscopy. Chemical Geology, 476, 248–259. 10.1016/j.chemgeo.2017.11.021

Thamdrup, B., Finster, K., Hansen, J. W., & Bak, F. (1993). Bacterial Disproportionation of Elemental Sulfur Coupled to Chemical Reduction of Iron or Manganese. Applied and Environmental Microbiology, 59(1), 101–108. 10.1128/aem.59.1.101-108.1993

ThomasArrigo, L. K., Bouchet, S., Kaegi, R., & Kretzschmar, R. (2020). Organic matter influences transformation products of ferrihydrite exposed to sulfide. Environmental Science: Nano, 7(11), 3405– 3418. 10.1039/d0en00398k

Truong, C., Bernard, S., Gorlas, A., & Guyot, F. (2024). Abiotic syntheses of pyrite: clues to assess the biogenicity of pyrite spherules. Geochemical Perspectives Letters, 32, 27–33. 10.7185/geochemlet.2438

Truong, C., Bernard, S., Le Pape, P., Morin, G., Baya, C., Merrot, P., Gorlas, A., & Guyot, F. (2023). Production of carbon-containing pyrite spherules induced by hyperthermophilic Thermococcales: a biosignature? Frontiers in Microbiology, 14, 1145781. 10.3389/fmicb.2023.1145781

Truong, C., Gaudu, N., Farr, O., Clouet, A., Ferry, D., Guyot, F., Ona-Nguema, G., Ruby, C., Nitschke, W., Grauby, O., & Duval, S. (2025). Fe2+ disproportionation within iron-rich alkaline vent analogues reveals proto-bioenergetic systems. Nature Communications, 16(1), 10682. 10.1038/s41467-025-65716-w

Tschech, A., & Pfennig, N. (1984). Growth yield increase linked to caffeate reduction in Acetobacterium woodii. Archives of Microbiology, 137, 163–167. 10.1007/BF00414460

van der Graaf, C. M., Sánchez-España, J., Ilin, A. M., Yusta, I., Stams, A. J. M., & Sánchez-Andrea, I. (2024). Micrometric pyrite catalyzes abiotic sulfidogenesis from elemental sulfur and hydrogen. Scientific Reports, 14(1), 17702. 10.1038/s41598-024-66006-z

Vigneron, A., Cruaud, P., Culley, A. I., Couture, R. M., Lovejoy, C., & Vincent, W. F. (2021). Genomic evidence for sulfur intermediates as new biogeochemical hubs in a model aquatic microbial ecosystem. Microbiome, 9, 46. 10.1186/s40168-021-00999-x

Višić, H., Duda, J.-P., Fischer, S., Escudero, C., Sekerci, F., Kappler, A., & Mansor, M. (2025). Experimental diagenesis reveals preservation of biosignatures in filamentous sulfur mats under hydrothermal conditions. Scientific Reports, 15, 38114. 10.1038/s41598-025-25172-4

Wan, M., Schröder, C., & Peiffer, S. (2017). Fe(III):S(-II) concentration ratio controls the pathway and the kinetics of pyrite formation during sulfidation of ferric hydroxides. Geochimica et Cosmochimica Acta, 217, 334–348. 10.1016/j.gca.2017.08.036

Wang, Q., & Morse, J. W. (1996). Pyrite formation under conditions approximating those in anoxic sediments I. Pathway and morphology. Marine Chemistry, 52, 99–121. 10.1016/0304-4203(95)00082-8

Wang, Q., Wang, J., Wang, X., Kumar; Naresh, Pan, Z., Peiffer, S., & Wang, Z. (2023). Transformations of Ferrihydrite–Extracellular Polymeric Substance Coprecipitates Driven by Dissolved Sulfide: Interrelated Effects of Carbon and Sulfur Loadings. Environ. Sci. Technol., 57(10), 4342–4353.

Williams, K. H., Ntarlagiannis, D., Slater, L. D., Dohnalkova, A., Hubbard, S. S., & Banfield, J. F. (2005). Geophysical imaging of stimulated microbial biomineralization. Environmental Science and Technology, 39(19), 7592–7600. 10.1021/es0504035

Wu, X.-T., Qiu, M., He, Y.-Q., Wu, K., Zhao, J.-Y., Wang, J., Ren, H.-Y., Su, J.-Y., & Bao, P. (2025). Disproportionation of elemental sulfur by Exiguobacterium from marine sediment. ISME Communications, 5(1), ycaf168. 10.1093/ismeco/ycaf168

Xiao, J., Hara, A. T., Kim, D., Zero, D. T., Koo, H., & Hwang, G. (2017). Biofilm three-dimensional architecture influences in situ pH distribution pattern on the human enamel surface. International Journal of Oral Science, 9, 74–79. 10.1038/ijos.2017.8

Yuan, S., Chou, I. M., Burruss, R. C., Wang, X., & Li, J. (2013). Disproportionation and thermochemical sulfate reduction reactions in S-H2O-CH4 and S-D2O-CH4 systems from 200 to 340°C at elevated pressures. Geochimica et Cosmochimica Acta, 118, 263–275. 10.1016/j.gca.2013.05.021

Zheng, S., Liao, M., Chen, Y., & Brook, M. A. (2020). Dissolving used rubber tires. Green Chemistry, 22(1), 94–102. 10.1039/c9gc03545a

Zhou, C., Vannela, R., Hayes, K. F., & Rittmann, B. E. (2014). Effect of growth conditions on microbial activity and iron-sulfide production by Desulfovibrio vulgaris. Journal of Hazardous Materials, 272, 28–35. 10.1016/j.jhazmat.2014.02.046

Zopfi, J., Ferdelman, T. G., & Fossing, H. (2004). Distribution and fate of sulfur intermediates - sulfite, tetrathionate, thiosulfate, and elemental sulfur - in marine sediments. In J. P. Amend, K. J. Edwards, & T. W. Lyons (Eds.), Sulfur Biogeochemistry - Past and Present (Vol. 379, pp. 97–116). Geological Society of America. 10.1130/0-8137-2379-5.97

