## Supplemental tables and figures for "A sulfur-cycling co-culture alters iron sulfide diagenesis and pyritization"

**Supplementary material**

| **Supplementary Table 1.** Summary of the follow-up experiments conducted to elucidate the presence of iron sulfides in sterile control incubations. | | | | |
| --- | --- | --- | --- | --- |
| Experiment | Round | Setup | Type of subsamples | Time of subsampling |
| Heating S_(0)_ at 30 and 80 °C | - | 2 sets of triplicate vials heated at 30 and 80 °C | • Liquid phase (formate, sulfate, thiosulfate) | T0, after 1 week, after 16 weeks |
|  |  |  | • Mineralogy | after 8 months |
| Abiotic sulfide production in the presence of ferrous iron | 1 | 2 sets of triplicate vials with either sulfate+ferrous iron or sulfate only. | • Liquid phase (lactate, sulfate), FeD | T0, after 4 weeks at 30 °C, after 16 weeks at 80 °C |
|  |  |  | • Mineralogy | once after 16 weeks at 80 °C |
|  | 2 | 2 sets of triplicate vials with either ferrous iron or sterile medium only (no sulfate or ferrous iron). | • Liquid phase (lactate, sulfate), FeD | once after 16 weeks at 80 °C |
|  |  |  | • Mineralogy | once after 16 weeks at 80 °C |

| **Supplementary Table 2.** Detailed mineralogical overview of the goethite experiment. | | | | | |
| --- | --- | --- | --- | --- | --- |
|  |  |  | Mineralogy | Morphology | Size |
| Incubation at 30 °C for four weeks | | | | | |
|  | Biotic incubation | Go + D. desulfuricans | goethite | acicular | ~ 1 μm |
|  |  |  | (oxidized) mackinawite, potentially greigite | aggregates of (nano)particles | < 1 μm |
|  |  | Go + S. deleyianum | goethite | acicular | ~ 1 μm |
|  |  |  | (oxidized) mackinawite, greigite | aggregates of (nano)particles | < 1 μm |
|  |  |  | elemental S(0) | not observed |  |
|  |  | Go + co-culture | goethite | acicular | ~ 1 μm |
|  |  |  | (oxidized) mackinawite, potentially greigite | aggregates of (nano)particles | < 1 μm |
|  |  |  | pyrite | aggregates of (nano)particles | < 1 μm |
|  | Abiotic incubation | Go+SO_4_^2-^+sulfide | goethite | acicular | ~ 1 μm |
|  |  | Go+S_(0)_+sulfide | goethite | acicular | ~ 1 μm |
|  |  | Go+*SO_4_^2-^+sulfide | goethite | acicular | ~ 1 μm |
|  | Control | Go+SO_4_^2-^ | goethite | acicular | ~ 1 μm |
|  |  | Go+S_(0)_ | goethite | acicular | ~ 1 μm |
|  |  | Go+*SO_4_^2-^ | goethite | acicular | ~ 1 μm |
| Incubation at 80 °C for 15-16 weeks (experimental diagenesis) | | | | | |
|  | Biotic incubation | Go + D. desulfuricans | goethite | acicular | ~ 1 μm |
|  |  |  | pyrite | euhedral/cubic, sometimes framboid-like | 1-2 μm |
|  |  |  | pyrite | spherules | 1-2 μm |
|  |  |  | vivianite | not observed |  |
|  |  | Go + S. deleyianum | goethite | acicular | ~ 1 μm |
|  |  |  | pyrite, greigite | aggregates of small crystals (cubes, irregular shapes) | ≤ 1 μm |
|  |  |  | pyrite (most likely) | euhedral/cubic | 1-2 μm |
|  |  |  | vivianite | not observed |  |
|  |  | Go + co-culture | goethite | acicular | ~ 1 μm |
|  |  |  | pyrite, marcasite | aggregates of small crystals (irregular shapes) | << 1 μm |
|  |  |  | pyrite (most likely) | euhedral/cubic | 1-2 μm |
|  |  |  | pyrite (most likely) | spherules | 1-2 μm |
|  |  |  | vivianite | not observed |  |
|  | Abiotic incubation | Go+SO_4_^2-^+sulfide | goethite | acicular | ~ 1 μm |
|  |  | Go+S_(0)_+sulfide | goethite | acicular | ~ 1 μm |
|  |  |  | vivianite | not observed |  |
|  |  | Go+*SO_4_^2-^+sulfide | goethite | acicular | ~ 1 μm |
|  | Control | Go+SO_4_^2-^ | goethite | acicular | ~ 1 μm |
|  |  | Go+S_(0)_ | goethite | acicular | ~ 1 μm |
|  |  |  | elemental S(0) | not observed |  |
|  |  | Go+*SO_4_^2-^ | goethite | acicular | ~ 1 μm |

| **Supplementary Table 3.** Detailed mineralogical overview of the ferric phosphate experiment. | | | | | |
| --- | --- | --- | --- | --- | --- |
|  |  |  | Mineralogy | Morphology | Size |
| Incubation at 30 °C for four weeks | | | | | |
|  | Biotic incubation | FP + D. desulfuricans | (oxidized) mackinawite, pyrite | sulfidated colloform aggregates | < 1 μm |
|  |  |  | vivianite | not observed |  |
|  |  | FP + S. deleyianum | (oxidized) mackinawite, pyrite | aggregates of (nano)particles | < 1 μm |
|  |  |  | vivianite | crystals with well-defined edges | > 10 μm |
|  |  |  | elemental S(0) | not observed |  |
|  |  | FP + co-culture | (oxidized) mackinawite | aggregates of (nano)particles | < 1 μm |
|  |  |  | vivianite | not observed |  |
|  | Abiotic incubation | FP+SO42^-^+sulfide | (oxidized) mackinawite, pyrite | aggregates of (nano)particles | < 1 μm |
|  |  |  | vivianite | not observed |  |
|  |  |  | elemental S(0) | not observed |  |
|  |  | FP+S_(0)_+sulfide | (oxidized) mackinawite, pyrite | aggregates of (nano)particles | < 1 μm |
|  |  |  | vivianite | not observed |  |
|  |  |  | elemental S(0) | not observed |  |
|  |  | FP+*SO_4_^2-^+sulfide | (oxidized) mackinawite, pyrite | aggregates of (nano)particles | < 1 μm |
|  |  |  | elemental S(0) | not observed |  |
|  | Control | FP+SO_4_^2-^ | ferric phosphate | colloform aggregate | < 1 μm |
|  |  | FP+S_(0)_ | ferric phosphate | colloform aggregate | < 1 μm |
|  |  |  | elemental S(0) | not observed |  |
|  |  | FP+*SO_4_^2-^ | ferric phosphate | colloform aggregate | < 1 μm |
| Incubation at 80 °C for 15-16 weeks (experimental diagenesis) | | | | | |
|  | Biotic incubation | FP + D. desulfuricans | pyrite | aggregates of spherules | ~ 1-2 μm |
|  |  |  | vivianite | prismatic or tabular crystals | > 10 μm |
|  |  | FP + S. deleyianum | pyrite | aggregates of spherules | ~ 1-2 μm |
|  |  |  | vivianite | prismatic or tabular crystals | > 10 μm |
|  |  | FP + co-culture | greigite, potentially pyrite | aggregates of small crystals (cubes, spherules or irregular shapes) | < 1 μm |
|  |  |  | vivianite | prismatic or tabular crystals | > 10 μm |
|  | Abiotic incubation | FP+SO42^-^+sulfide | pyrite | euhedral/cubic, sometimes framboid-like | ~ 1-2 μm |
|  |  |  | vivianite | prismatic or tabular crystals | > 10 μm |
|  |  | FP+S_(0)_+sulfide | pyrite | aggregates of spherules | ~ 1-2 μm |
|  |  |  | vivianite | prismatic or tabular crystals | > 10 μm |
|  |  | FP+*SO_4_^2-^+sulfide | pyrite | aggregates of spherules | ~ 1-2 μm |
|  |  |  | vivianite | prismatic or tabular crystals | > 10 μm |
|  | Control | FP+SO_4_^2-^ | phosphate mineral/ferric phosphate | flower-like structures | 1-2 μm |
|  |  | FP+S_(0)_ | pyrite | aggregates of spherules | ~ 1-2 μm |
|  |  |  | phosphate mineral/ferric phosphate | flower-like structures | 1-2 μm |
|  |  | FP+*SO_4_^2-^ | phosphate mineral/ferric phosphate | flower-like structures | 1-2 μm |

| **Supplementary Table 4.** Detailed mineralogical overview of the ferrihydrite experiment. | | | | | |
| --- | --- | --- | --- | --- | --- |
|  |  |  | Mineralogy | Morphology | Size |
| Incubation at 30 °C for four weeks | | | | | |
|  | Biotic incubation | Fh + D. desulfuricans | No growth | No growth |  |
|  |  | Fh + S. deleyianum | (oxidized) mackinawite | aggregates of (nano)particles | < 1 μm |
|  |  |  | elemental S(0) | not observed |  |
|  |  | Fh + co-culture | (oxidized) mackinawite | aggregates of (nano)particles | < 1 μm |
|  |  |  | (oxidized) mackinawite | flat sheets | ~ 10 μm |
|  | Abiotic incubation | Fh+SO_4_^2-^+sulfide | (oxidized) mackinawite | aggregates of (nano)particles | < 1 μm |
|  |  | Fh+S_(0)_+sulfide | (oxidized) mackinawite | aggregates of (nano)particles | < 1 μm |
|  |  |  | vivianite | not observed |  |
|  |  |  | elemental S(0) | not observed |  |
|  |  | Fh+*SO_4_^2-^+sulfide | (oxidized) mackinawite | aggregates of (nano)particles | < 1 μm |
|  |  |  | elemental S(0) | not observed |  |
|  | Control | Fh+SO_4_^2-^ | ferrihydrite | aggregates of (nano)crystals | < 1 - 10 μm |
|  |  | Fh+S_(0)_ | ferrihydrite | aggregates of (nano)crystals | < 1 - 10 μm |
|  |  | Fh+*SO_4_^2-^ | ferrihydrite | aggregates of (nano)crystals | < 1 - 10 μm |
| Incubation at 80 °C for 15-16 weeks (experimental diagenesis | | | | | |
|  |  |  | Mineralogy | Morphology | Size |
|  | Biotic incubation | Fh + D. desulfuricans | No growth | No growth |  |
|  |  | Fh + S. deleyianum | (oxidized) mackinawite | flat sheets | ~ 10 μm |
|  |  |  | pyrite | aggregates of spherules | ~ 1-2 μm |
|  |  | Fh + co-culture | (oxidized) mackinawite | flat sheets | ~ 10 μm |
|  |  |  | greigite, pyrite | aggregates of spherules | ~ 1-2 μm |
|  |  |  | vivianite | not observed |  |
|  | Abiotic incubation | Fh+SO_4_^2-^+sulfide | (oxidized) mackinawite | flat sheets | ~ 10 μm |
|  |  |  | pyrite | euhedral/octahedral crystals | ~ 1-2 μm |
|  |  | Fh+S_(0)_+sulfide | (oxidized) mackinawite | flat sheets | ~ 10 μm |
|  |  |  | pyrite | aggregates of spherules | ~ 1-2 μm |
|  |  |  | vivianite | not observed |  |
|  |  | Fh+*SO_4_^2-^+sulfide | (oxidized) mackinawite | flat sheets | ~ 10 μm |
|  |  |  | pyrite (most likely) | euhedral/cubic/octahedral crystals | ~ 1-2 μm |
|  |  |  | pyrite (most likely) | aggregates of spherules | ~ 1-2 μm |
|  | Control | Fh+SO_4_^2-^ | ferrihydrite | aggregates of (nano)crystals | < 1 - 10 μm |
|  |  | Fh+S_(0)_ | pyrite | aggregates of spherules | ~ 1-2 μm |
|  |  |  | ferrihydrite | aggregates of (nano)crystals | < 1 - 10 μm |
|  |  |  | elemental S(0) | not observed |  |
|  |  | Fh+*SO_4_^2-^ | ferrihydrite | aggregates of (nano)crystals | < 1 - 10 μm |

| **Supplementary Table 5.** Detailed mineralogical overview of the ferrous iron experiment. | | | | | |
| --- | --- | --- | --- | --- | --- |
|  |  |  | Mineralogy | Morphology | Size |
| Incubation at 30 °C for four weeks | | | | | |
|  | Biotic incubation | Fe^2+^ + D. desulfuricans | mackinawite, potentially greigite | aggregates of (nano)particles | < 1 μm |
|  |  | Fe^2+^ + S. deleyianum | mackinawite, greigite | aggregates of (nano)particles | < 1 μm |
|  |  |  | vivianite | not observed |  |
|  |  |  | elemental S(0) | not observed |  |
|  |  | Fe^2+^ + co-culture | mackinawite, potentially greigite | aggregates of (nano)particles | < 1 μm |
|  |  |  | elemental S(0) | not observed |  |
|  | Abiotic incubation | Fe^2+^+SO_4_^2-^+sulfide | mackinawite | aggregates of (nano)particles | < 1 μm |
|  |  | Fe^2+^+S_(0)_+sulfide | mackinawite, potentially greigite | aggregates of (nano)particles | < 1 μm |
|  |  |  | elemental S(0) | not observed |  |
|  |  | Fe^2+^+SO_4_^2-^/S_(0)_+sulfide | mackinawite, potentially greigite | aggregates of (nano)particles | < 1 μm |
|  | Control | Fe^2+^+SO_4_^2-^ | No sample |  |  |
|  |  | Fe^2+^+S_(0)_ | No sample |  |  |
|  |  | Fe^2+^+SO_4_^2-^/S_(0)_ | No sample |  |  |
| Incubation at 80 °C for 15-16 weeks (experimental diagenesis) | | | |  |  |
|  | Biotic incubation | Fe^2+^ + D. desulfuricans | mackinawite, greigite, pyrite | aggregates of (nano)particules that often form larger spherules | < 1 μm |
|  |  | Fe^2+^ + S. deleyianum | (oxidized) mackinawite, pyrite | aggregates of (nano)particules that often form larger spherules | < 1 μm |
|  |  | Fe^2+^ + co-culture | greigite (mostly), pyrite | aggregates of spherules | < 1 μm or 2-5 μm |
|  |  |  | elemental S(0) | not observed |  |
|  | Abiotic incubation | Fe^2+^+SO_4_^2-^+sulfide | mackinawite | aggregates of (nano)particles | < 1 μm |
|  |  |  | pyrite | euhedral/cubic crystals | ~ 1-2 μm |
|  |  | Fe^2+^+S_(0)_+sulfide | (oxidized) mackinawite, pyrite | aggregates of spherulitic (nano)particles | < 1 μm |
|  |  | Fe^2+^+SO_4_^2-^/S_(0)_+sulfide | greigite, pyrite | aggregates of spherulitic (nano)particles | < 1 μm |
|  |  |  | elemental S(0) | not observed |  |
|  | Control | Fe^2+^+SO_4_^2-^ | (oxidized) mackinawite, pyrite, marcasite, | structures that contain Fe and S (spherules, octahedra) | ~ 1-2 μm |
|  |  |  | iron phosphate mineral | spherules | ~ 1-5 μm |
|  |  |  | magnetite | structures that contain Fe and O (sheets, acicular shapes) | ~ 1-2 μm |
|  |  | Fe^2+^+S_(0)_ | pyrite | aggregates of spherulitic (nano)particles | < 1 μm |
|  |  |  | iron phosphate mineral | spherules | ~ 1-5 μm |
|  |  | Fe^2+^+SO_4_^2-^/S_(0)_ | pyrite | aggregates of spherulitic (nano)particles | < 1 μm |
|  |  |  | iron phosphate mineral | spherules | ~ 1-5 μm |

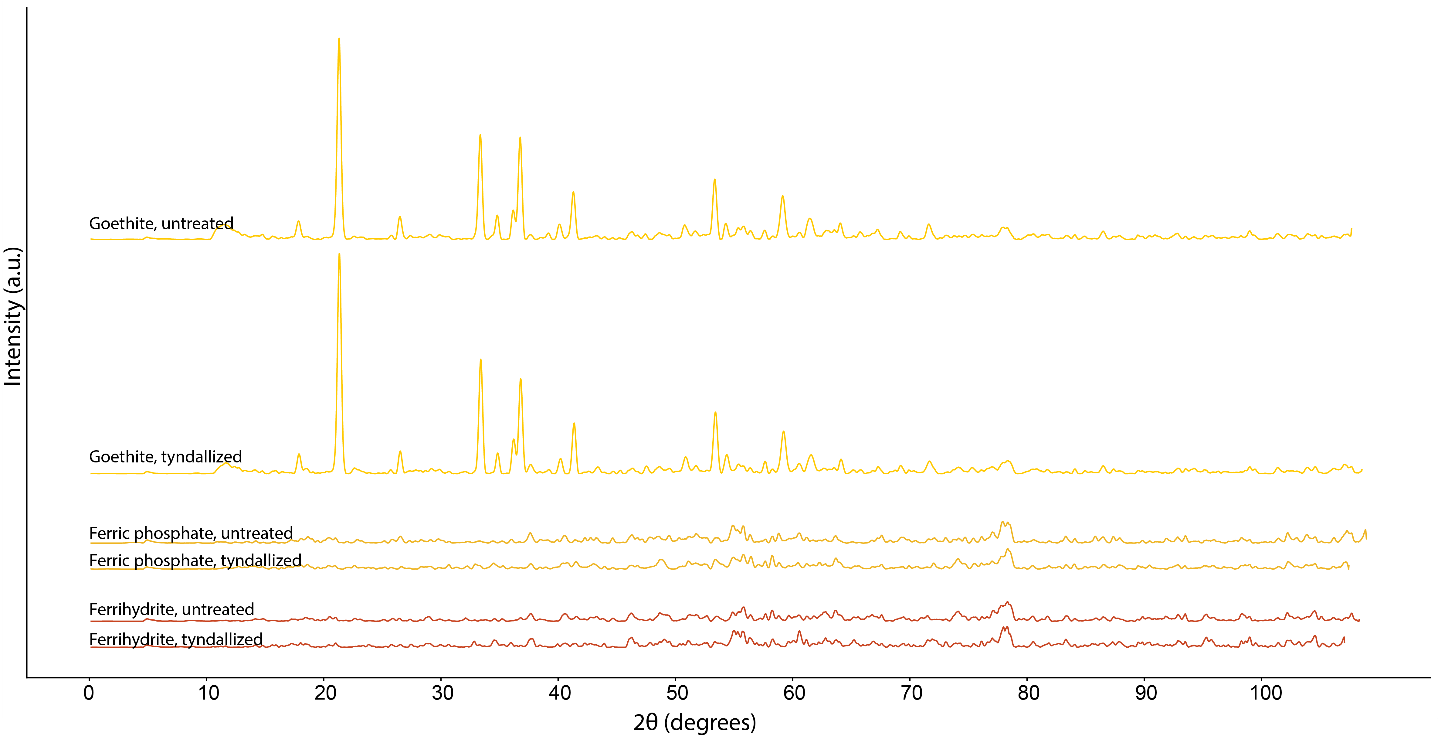

**Figure S1**. X-ray diffractograms of the untreated and tyndallized (one hour at 90 °C, repeated on three consecutive days) ferric compounds.

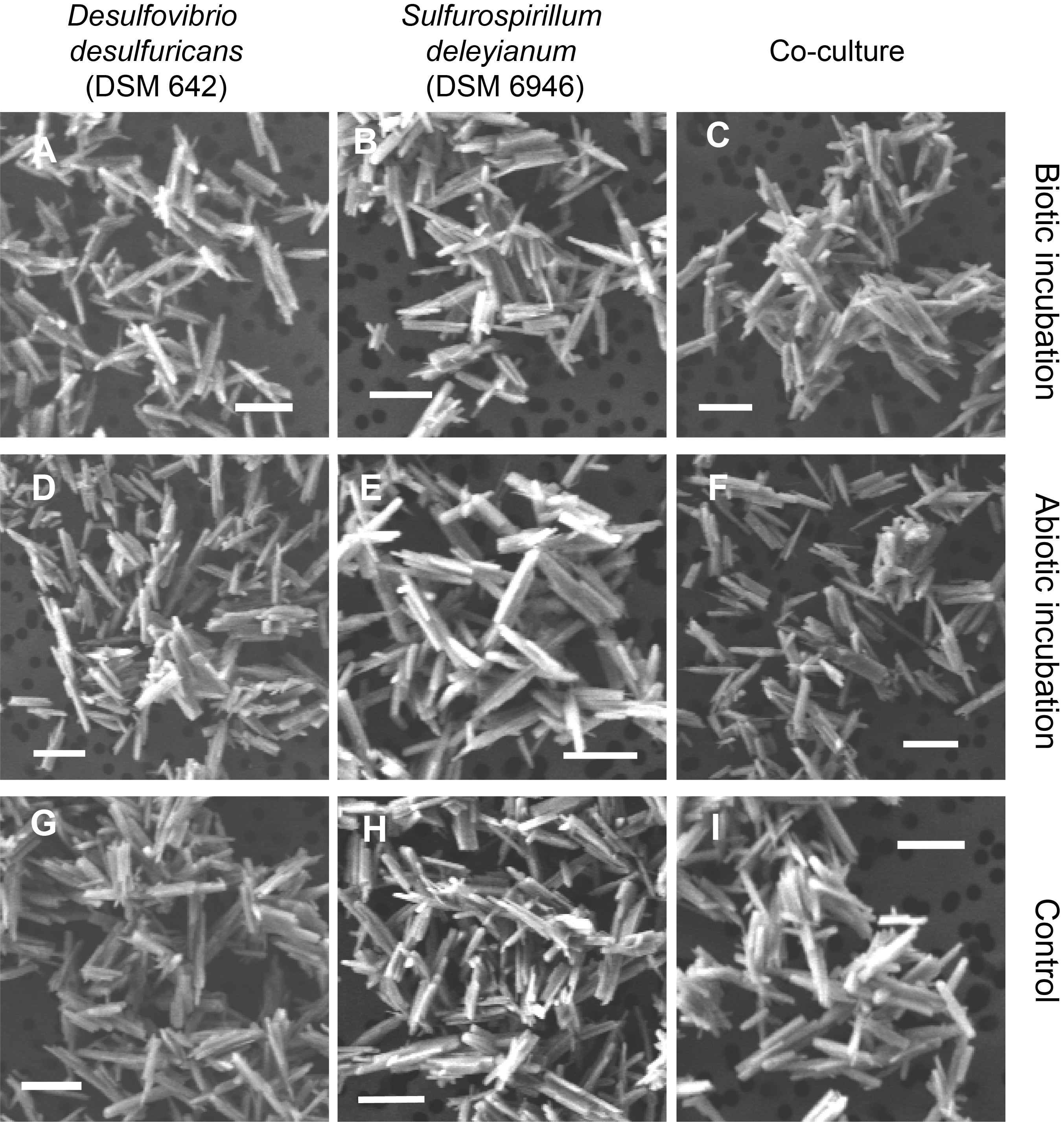

**Figure S2**. SEM images of the goethite T_0_ samples. All images were taken in secondary electron (SE) mode.

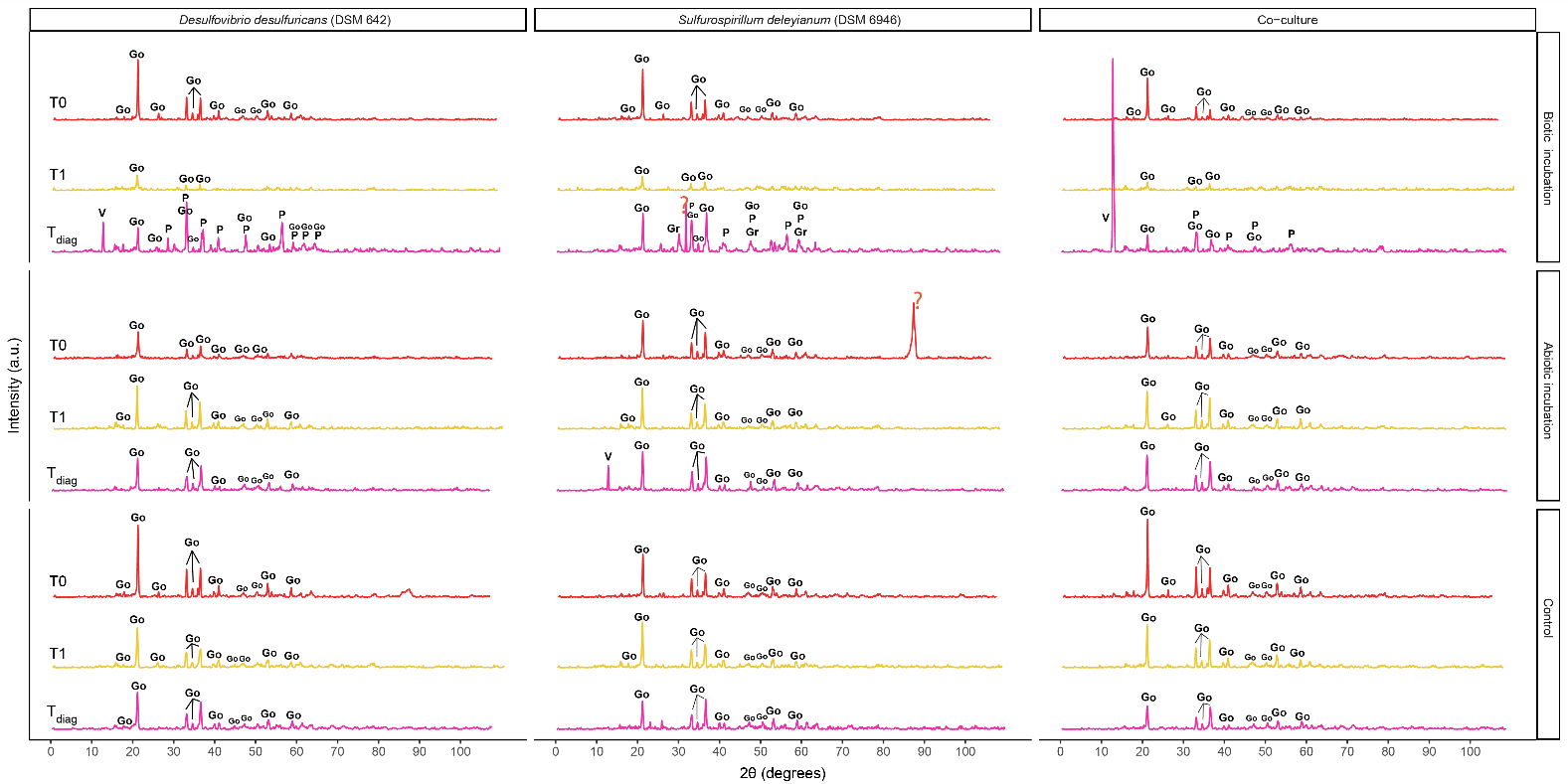

**Figure S3**. X-ray diffractograms of the goethite biomineralization experiment. Abbreviations: T_0_ – subsamples taken at the start of the experiments, T1– subsamples taken after 1 month at 30 °C, T_diag_ – subsamples taken after 15-16 weeks of experimental diagenesis (80 °C), P – pyrite, Gr – greigite, Go – goethite, V – vivianite.

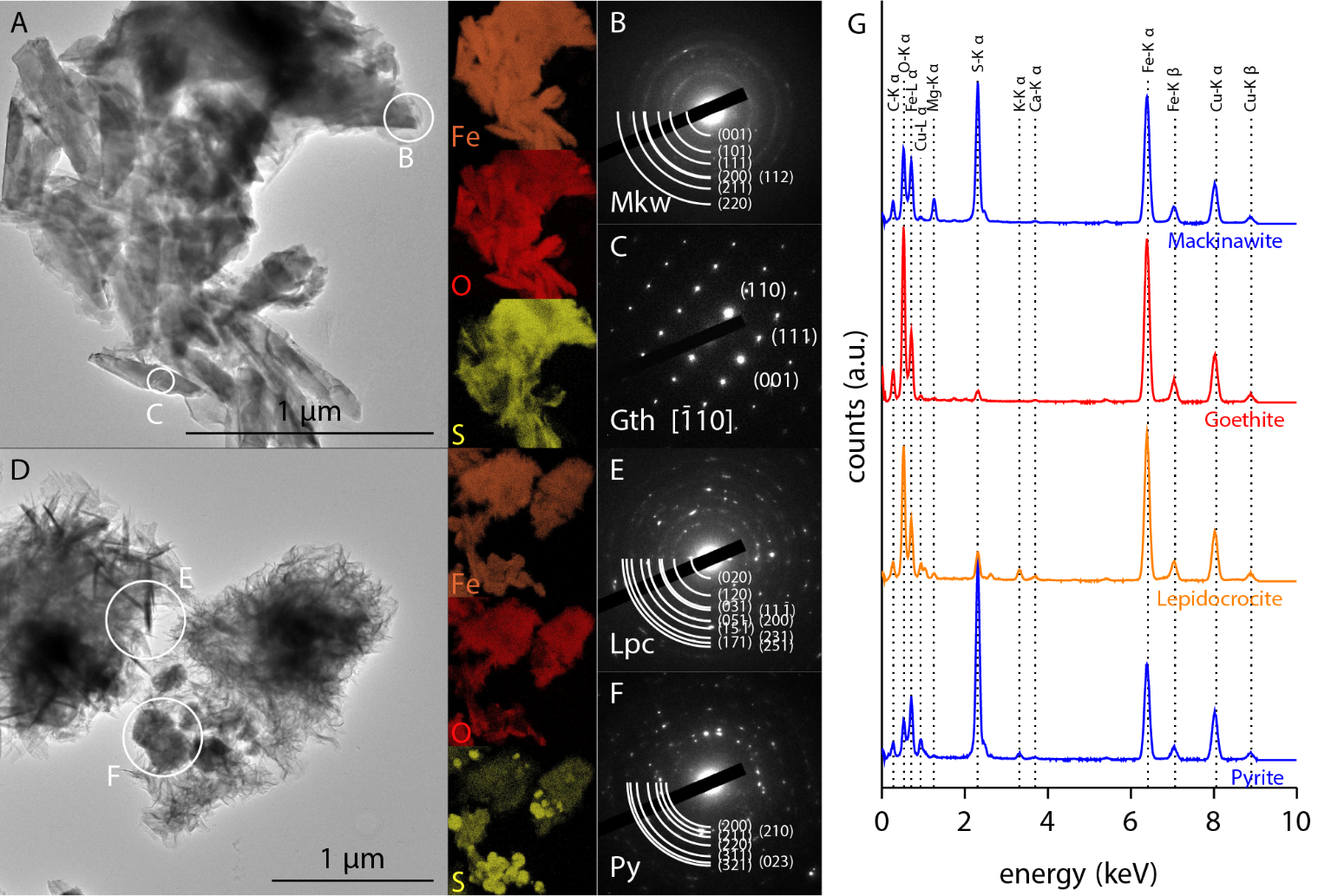

**Figure S4.** TEM data of the goethite biomineralization experiments. A –TEM image and associated Fe, O and S EDS mapping of produced mackinawite and residual goethite in the S. deleyianum culture. B – SAED pattern of poorly crystallized mackinawite. C – SAED pattern of goethite monocrystal. D – TEM image and associated Fe, O and S EDS mapping of lepidocrocite and pyrite produced in the co-culture incubation. E – SAED pattern of polycrystalline lepidocrocite. F – SAED pattern of polycrystalline pyrite. G – EDS spectra of regions of interest corresponding to the SAEDs.

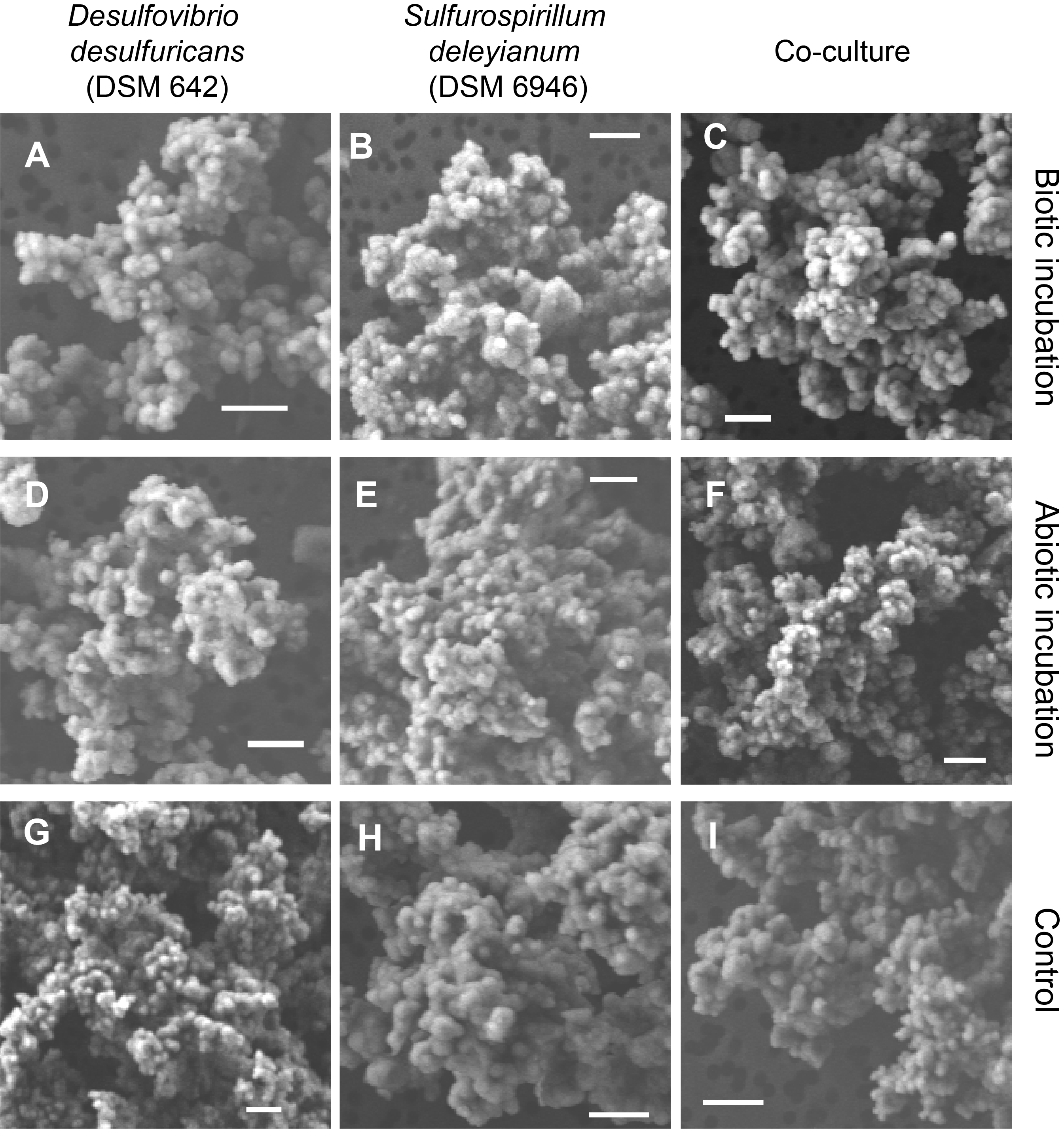

**Figure S5**. SEM images of the ferric phosphate T_0_ samples. All images were taken in secondary electron (SE) mode.

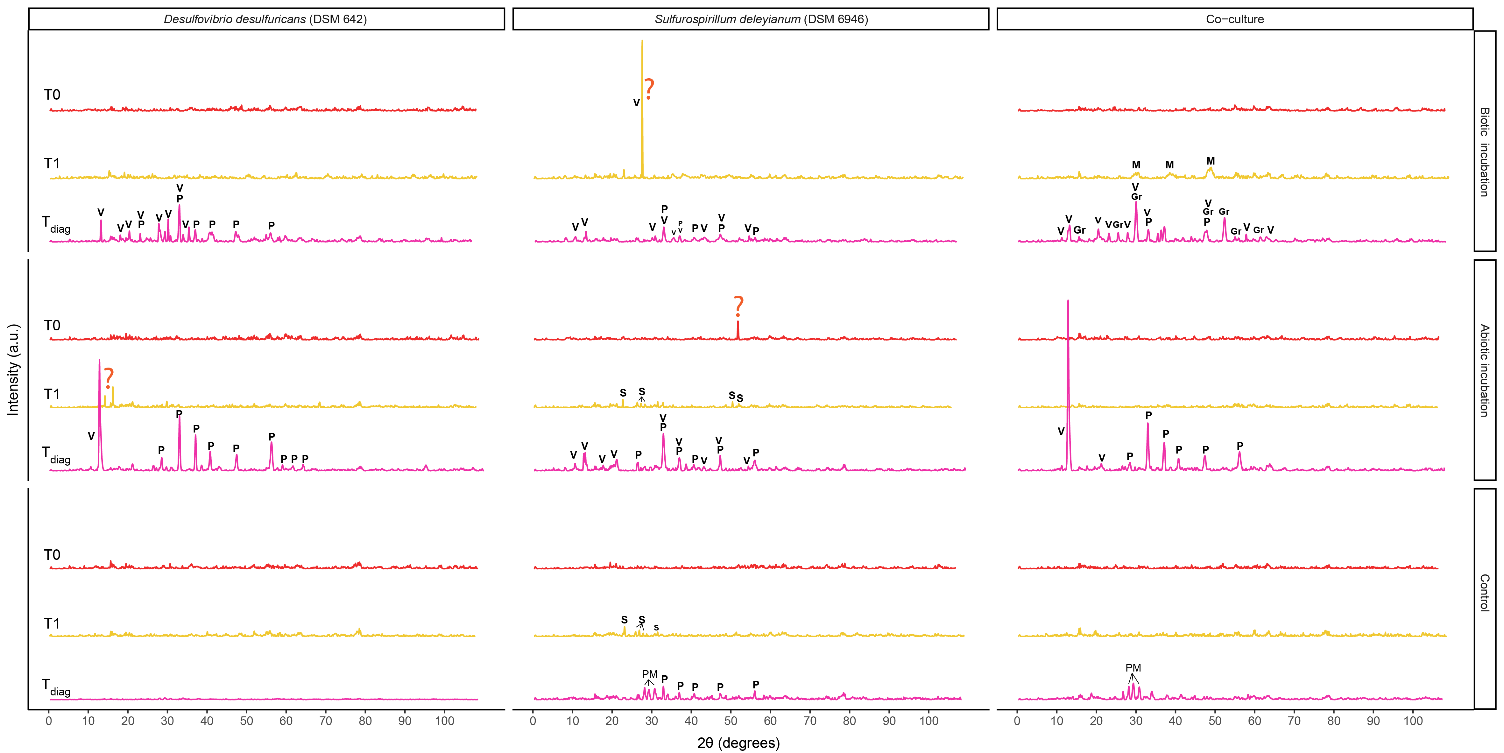

**Figure S6.** X-ray diffractograms of the ferric phosphate biomineralization experiment. Abbreviations: T_0_ – subsamples taken at the start of the experiments, T1– subsamples taken after 1 month at 30 °C, T_diag_ – subsamples taken after 15-16 weeks of experimental diagenesis (80 °C), P – pyrite, M – mackinawite, Gr – greigite, V – vivianite, PM – phosphate mineral.

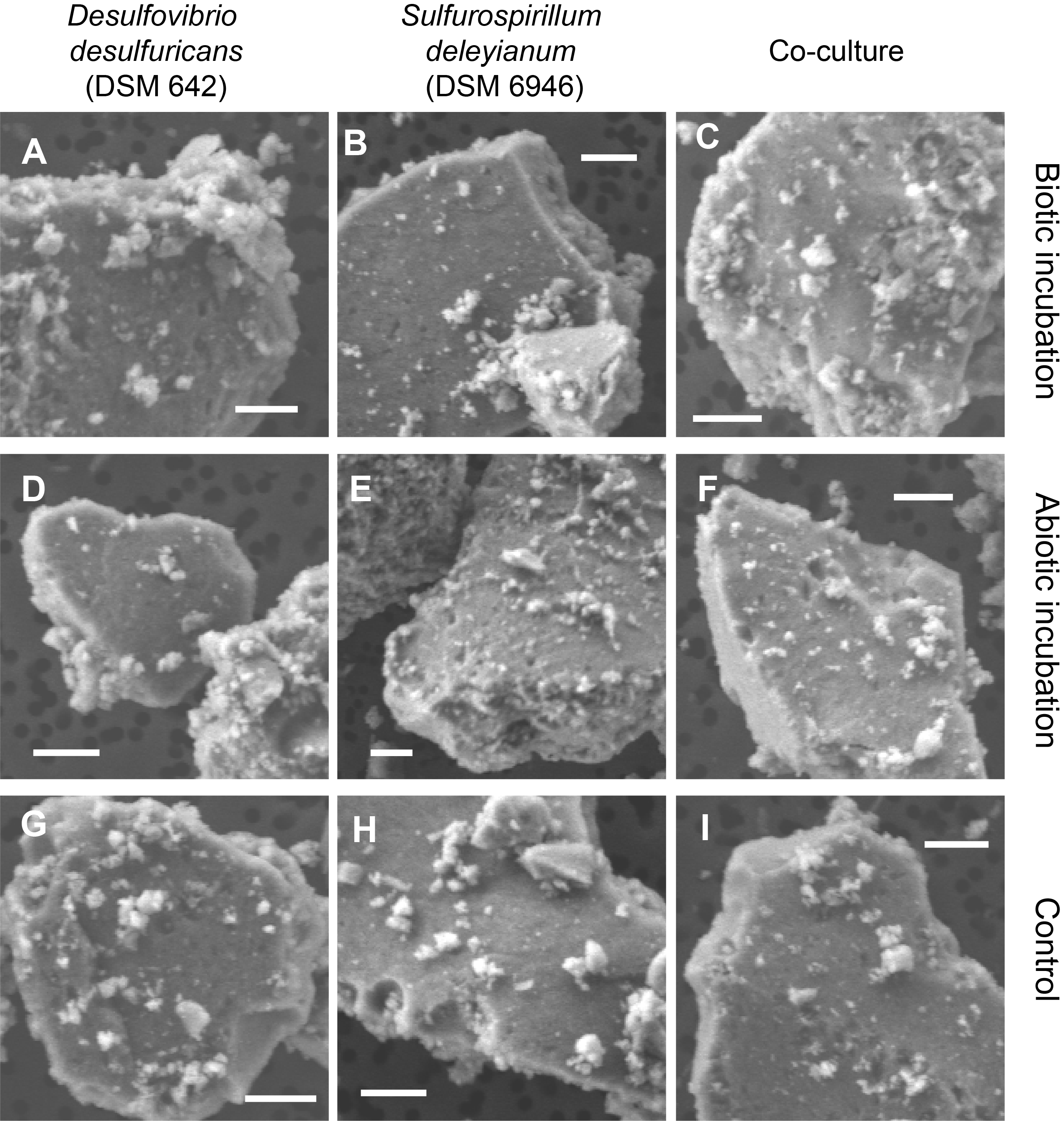

**Figure S7**. SEM images of the ferrihydrite T_0_ samples. All images were taken in secondary electron (SE) mode.

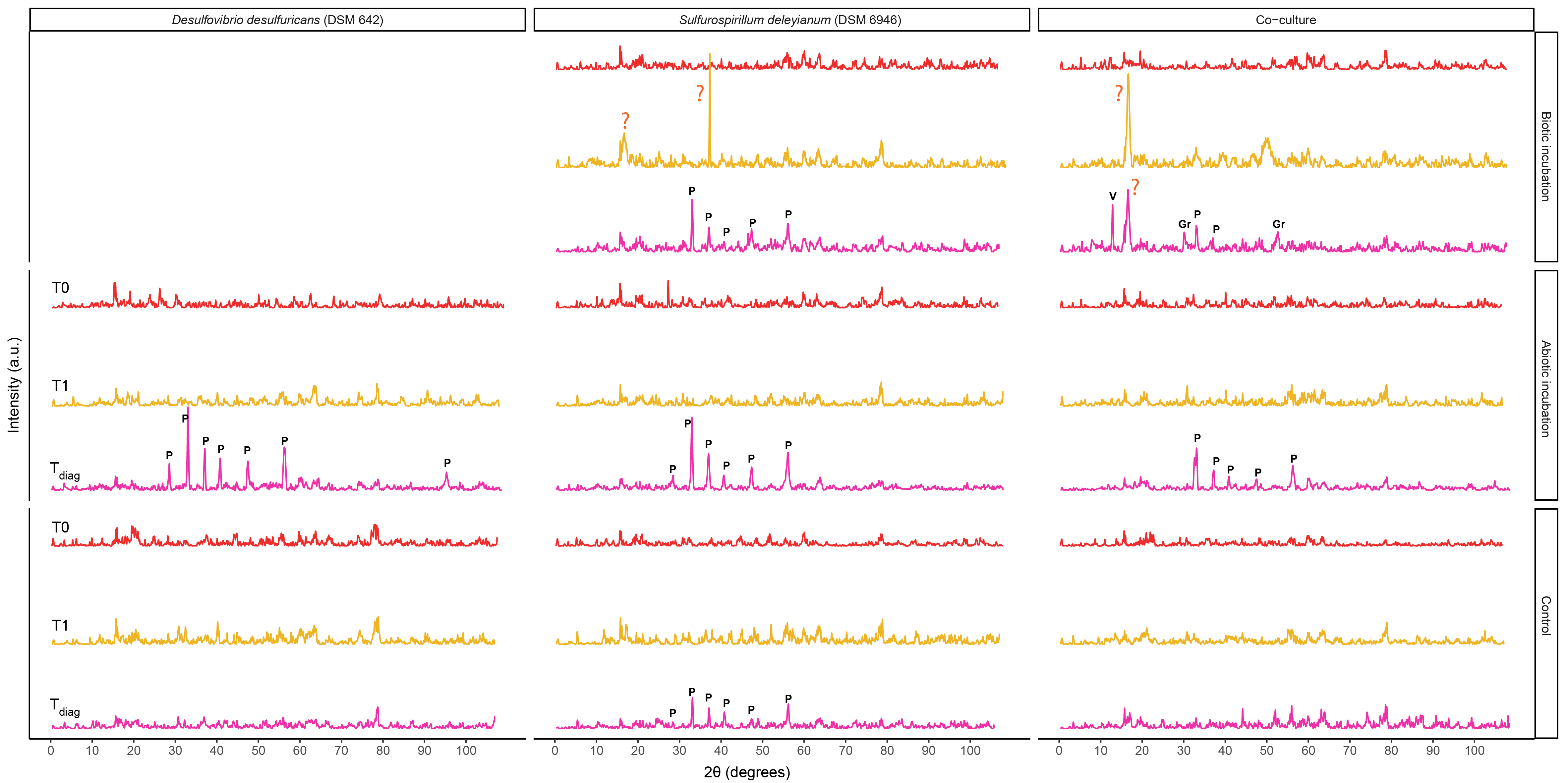

**Figure S8.** X-ray diffractograms of the ferrihydrite biomineralization experiment. Abbreviations: T_0_ – subsamples taken at the start of the experiments, T1– subsamples taken after 1 month at 30 °C, T_diag_ – subsamples taken after 15-16 weeks of experimental diagenesis (80 °C), P – pyrite, Gr – greigite, V – vivianite.

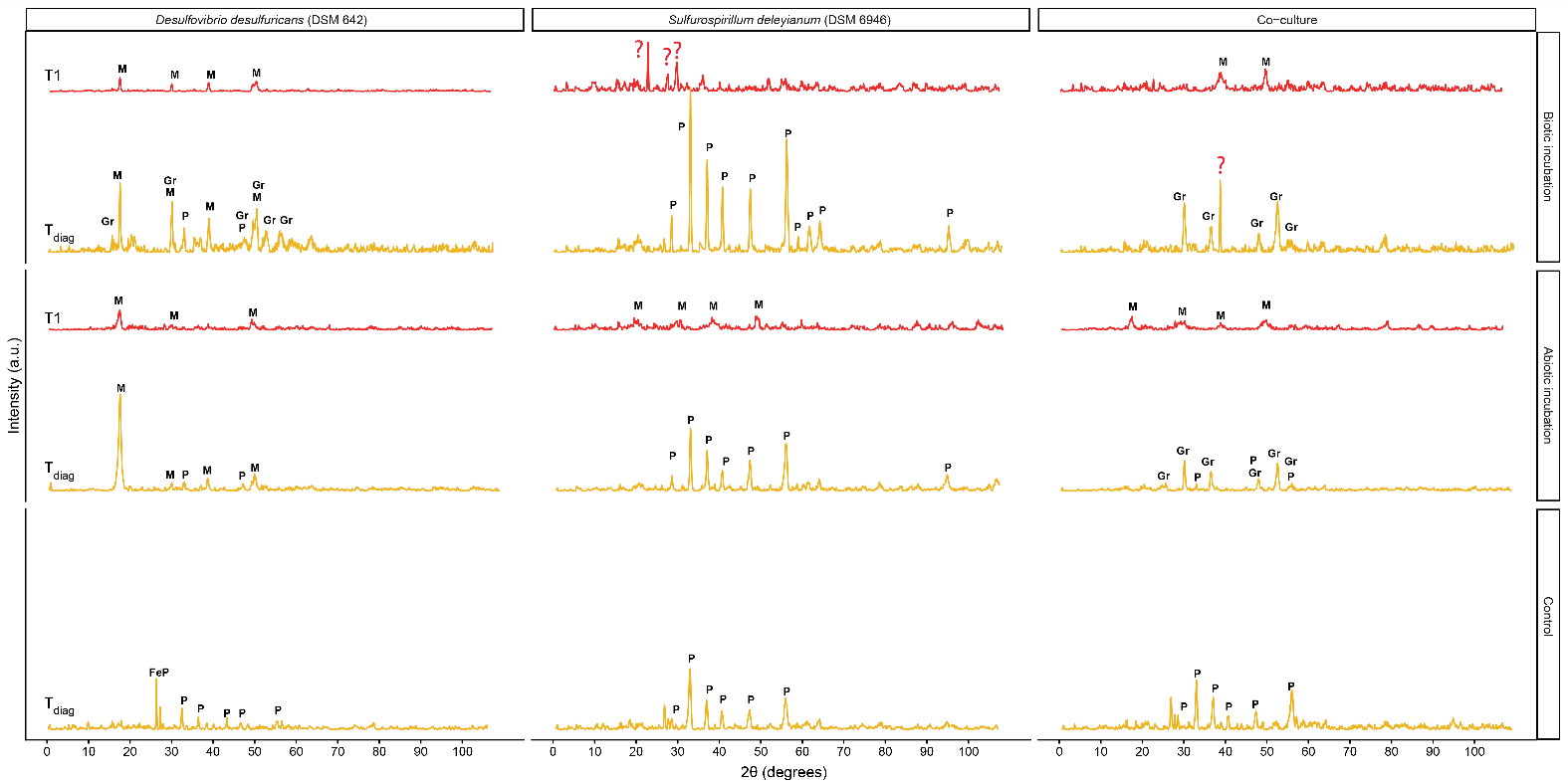

**Figure S9.** X-ray diffractograms of the ferrous chloride biomineralization experiment. Abbreviations: T_0_ – subsamples taken at the start of the experiments, T1– subsamples taken after 1 month at 30 °C, T_diag_ – subsamples taken after 15-16 weeks of experimental diagenesis (80 °C), P – pyrite, M – mackinawite, Gr – greigite, FeP – iron phosphate.

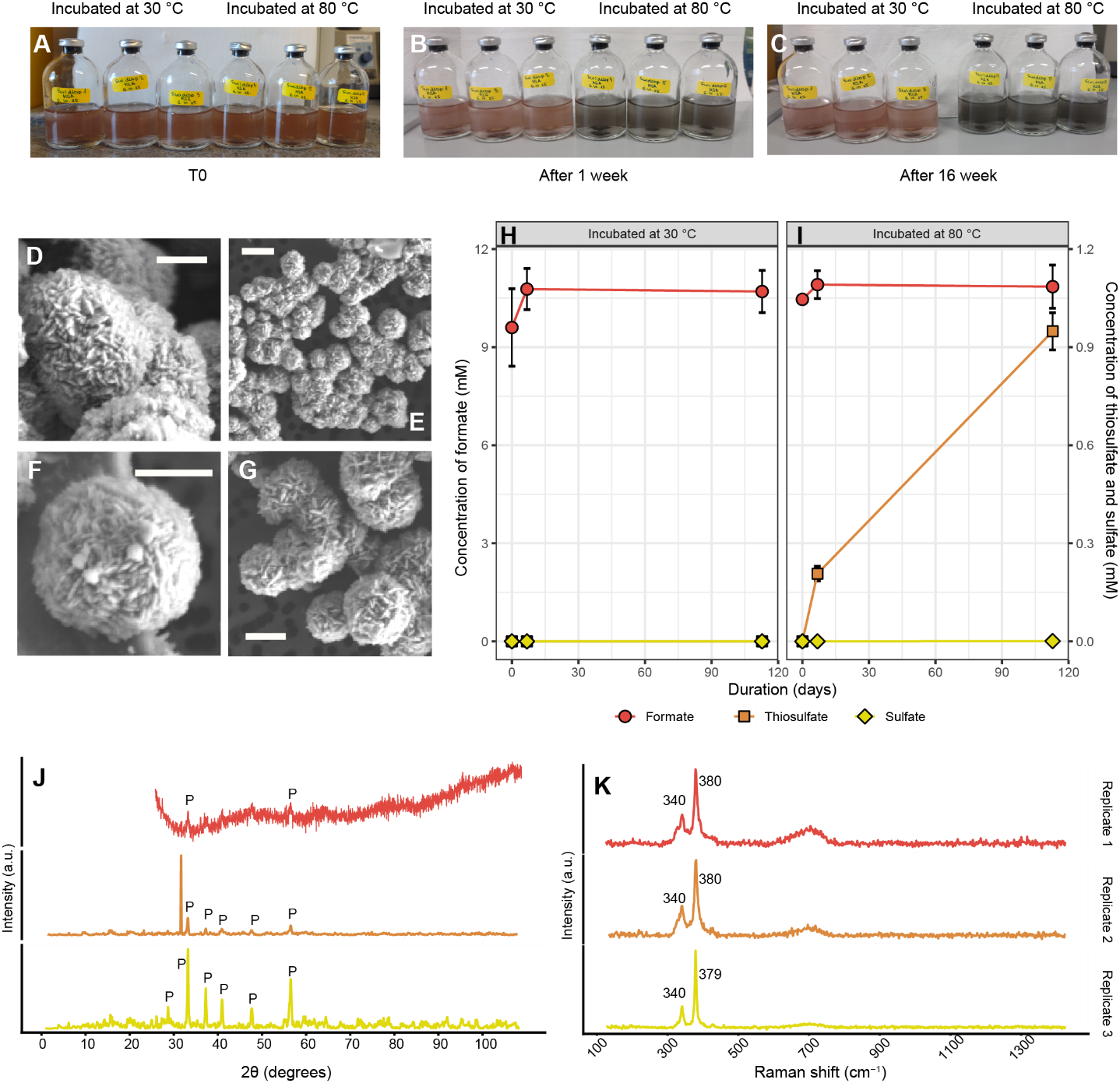

**Figure S10.** The effect of heating elemental S_(0)_ (10 mM) in the presence of ferrihydrite (8 mM) at 30 and 80 °C. A, B and C – physical appearance of serum vials at the start of the experiment (T0), after 1 week and after 16 weeks, respectively. D, E, F, and G – SEM images of pyrite spherules incubated for 8 months at 80 °C. H – Concentration of sulfate, thiosulfate and formate in vials incubated at 30 °C. I – Concentration of sulfate, thiosulfate and formate in vials incubated at 80 °C. J – X-ray diffractograms of the three replicates incubated for 8 months at 80 °C. Abbreviations: P – pyrite. In Replicate 1, no background removal was done. K – Raman spectroscopy results of the three replicates incubated for 8 months at 80 °C. Error bars represent the standard deviation from the average of three replicates.

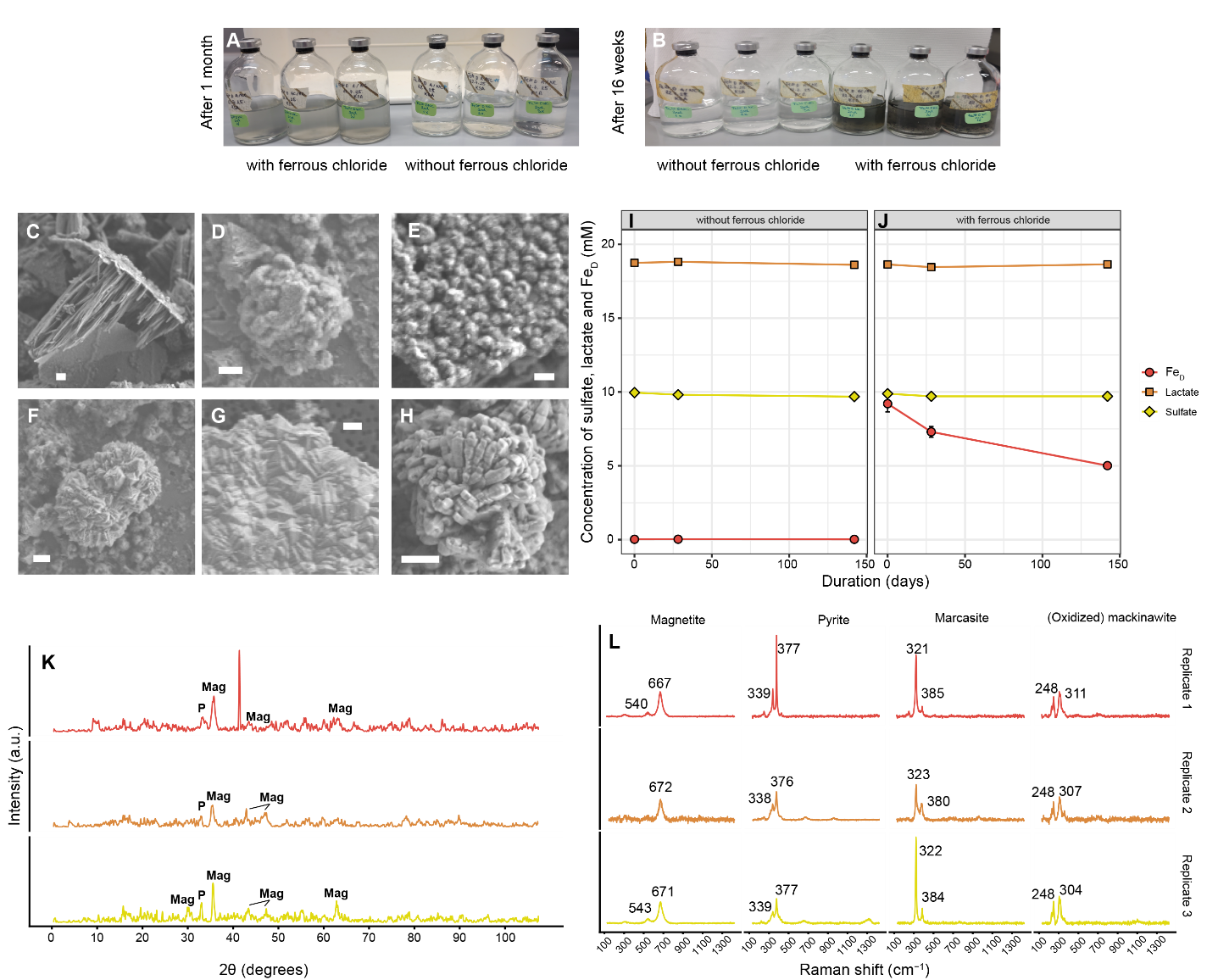

**Figure S11.** The effect of heating sulfate in the presence of ferrous iron at 30 and 80 °C. A, B – physical appearance of serum vials after 1 month at 30 °C and after 16 weeks at 80 °C, respectively. C, D, E, F, G and H – SEM images of various structures containing Fe, S and O (see text). I – Concentration of sulfate, lactate and dissolved iron (Fe_D_) in vials without added ferrous chloride. J – Concentration of sulfate, lactate and dissolved iron (Fe_D_) in vials with added ferrous chloride. K – X-ray diffractograms of the three replicates. Abbreviations: P – pyrite, Mag – magnetite. L – Raman spectroscopy results of the three replicates. Error bars represent the standard deviation from the average of three replicates.

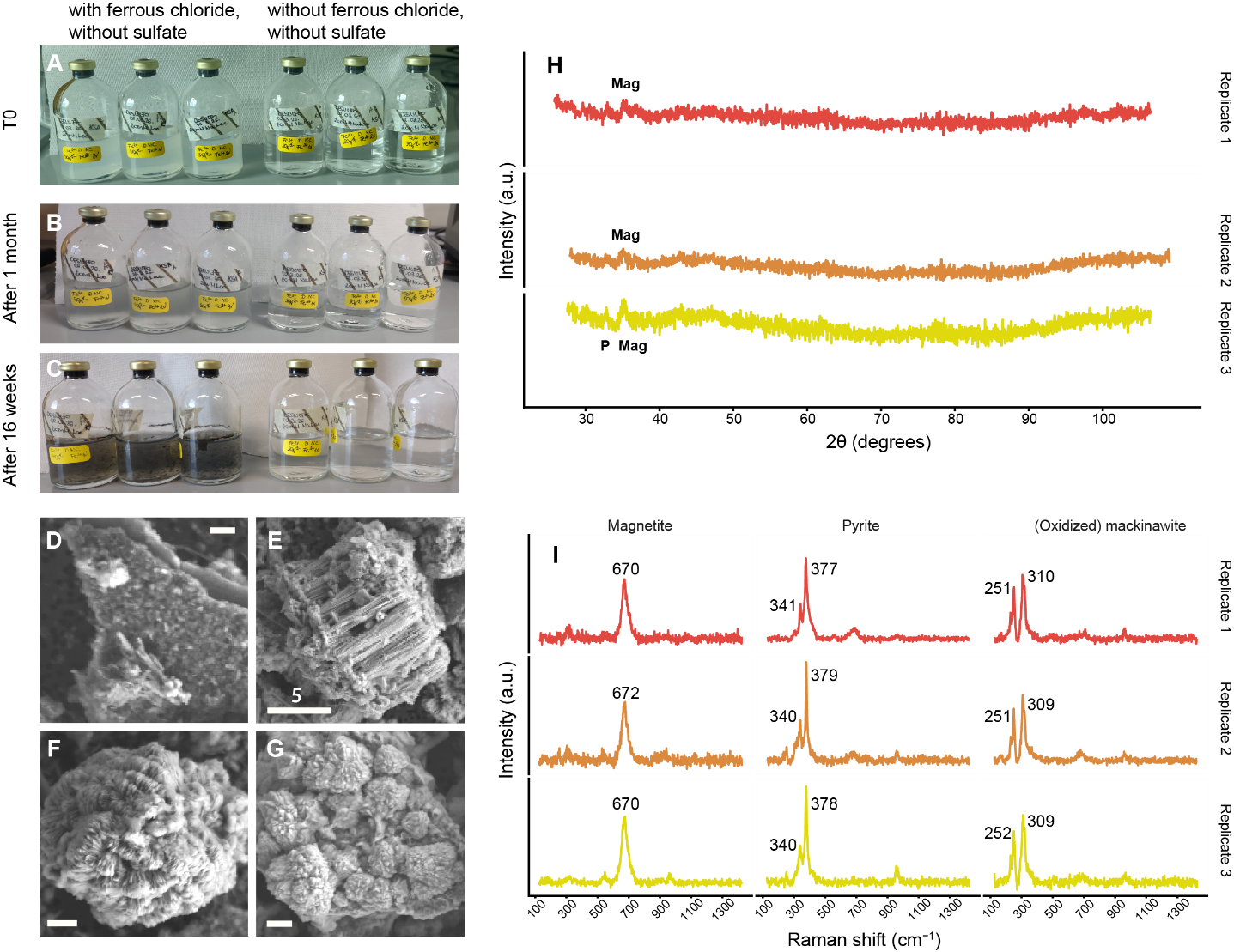

**Figure S12**. The effect of heating the sterile cultivation medium without sulfate in the presence of ferrous iron at 30 and 80 °C. A, B and C – physical appearance of serum vials at the start of the experiment (T0), after 1 month at 30 °C and after 16 weeks at 80 °C, respectively. D, E, F and G – SEM images of various structures containing Fe, S and O (see text). H – X-ray diffractograms of the three replicates. Abbreviations: P – pyrite, Mag – magnetite. I – Raman spectroscopy results of the three replicates.
